# Longitudinal analysis of memory T follicular helper cells and antibody response following CoronaVac vaccination

**DOI:** 10.1101/2023.05.16.541033

**Authors:** Pengcheng Zhou, Cheng Cao, Tuo Ji, Ting Zheng, Yaping Dai, Min Liu, Junfeng Jiang, Daoqi Sun, Zhonghu Bai, Xiaojie Lu, Fang Gong

**Affiliations:** Department of Laboratory Medicine, Affiliated Hospital of Jiangnan University, Wuxi, Jiangsu, China; The University of Queensland Diamantina Institute, Brisbane, Australia; Wuxi School of Medicine, Jiangnan University, Wuxi, Jiangsu, China; Department of central laboratory, The Second People’s Hospital of Lianyungang City (Cancer Hospital of Lianyungang), Lianyungang, Jiangsu, China; Hospital for Special Surgery, Weill Cornell Medical College, New York, NY, USA; Department of Laboratory Medicine, The Fifth People’s Hospital of Wuxi, Wuxi, Jiangsu, China; School of Biotechnology, Jiangnan University, Wuxi, Jiangsu, China

## Abstract

The inactivated vaccine CoronaVac is one of the most widely used COVID-19 vaccines globally. However, the longitudinal evolution of the immune response induced by CoronaVac remains elusive compared to other vaccine platforms. Here, we recruited 88 healthy individuals that received 3 doses of CoronaVac vaccine. We longitudinally evaluated their polyclonal and antigen-specific CD4^+^ T cells and neutralizing antibody response after receiving each dose of vaccine for over 300 days. Both the 2^nd^ and 3^rd^ dose of vaccination induced robust spike-specific neutralizing antibodies, with a 3^rd^ vaccine further increased the overall magnitude of antibody response, and neutralization against Omicron sub-lineages B.1.1.529, BA.2, BA.4/BA.5 and BA.2.75.2. Spike-specific CD4^+^ T cell and circulating T follicular helper (cT_FH_) cells were markedly increased by the 2^nd^ and 3^rd^ dose of CoronaVac vaccine, accompanied with altered composition of functional cT_FH_ cell subsets with distinct effector and memory potential. Additionally, cT_FH_ cells are positively correlated with neutralizing antibody titers. Our results suggest that CoronaVac vaccine-induced spike-specific T cells are capable of supporting humoral immunity for long-term immune protection.

## Introduction

Since the onset of Coronavirus disease (COVID-19) pandemic in 2020, the SARS-CoV-2 virus has evolved into several sub-lineages, including Alpha, Beta, Gamma, Delta, and Omicron (1). Such continuous evolution resulted in rapid and divergent mutations over Omicron, with further Omicron sub-lineages such as B1.1.529, BA.2, BA.3, BA.4/5, BA.2.75, BQ1.1 currently spreading in many countries. Despite the significant success of human vaccine efforts in preventing severe disease caused by both ancestral and emerging strains of SARS-CoV-2, there are still significant knowledge gaps in our understanding of the mechanisms of vaccine-induced immune protection against emerging variants. These gaps are particularly evident in studies of different vaccine platforms, indicating an inequitable distribution of knowledge. Unlike mRNA vaccines, the evolution and persistence of human immune response elicited by vaccine platforms such as inactivated vaccines or protein vaccines after the second and third booster doses are poorly understood. Recent evidence has shown that a homologous 3^rd^ dose of CoronaVac was associated with an efficient increase in SARS-CoV-2-specific antibodies (2–4). However, the human immune response and antibody neutralizing capacity against ancestral and Omicron sub-lineages including BA.2, BA.4/5 and BA.2.75.2 by a 3^rd^ dose of CoronaVac vaccine remains largely unexplored.

In addition to the serologic antibodies, antigen-specific CD4^+^ T cells, especially T follicular helper cells (T_FH_) are critical for the long-term immune protection (5–7). mRNA vaccines induced robust antigen-specific memory CD4^+^ T cells and T follicular helper cells (8, 9). However, the dynamics of CoronaVac-induced memory CD4^+^ T cells and their relationship with prolonged antibody response remains poorly understood. It is also crucial to determine whether certain subsets of human T_FH_ cells exhibit superior effector or memory potential in response to a vaccine longitudinally. Here, we recruited 88 healthy individuals in a longitudinal cohort who received three doses of inactivated CoronaVac vaccines. In ordered to understand the longevity and nature of human immune response induced by CoronaVac vaccine, we evaluated polyclonal and antigen-specific T_FH_ cells, neutralizing antibodies and their relationships following primary, secondary and third booster of CoronaVac vaccinations.

## Results

### Spike-specific antibody response elicited by CoronaVac vaccine over time

88 healthy individuals were recruited in this study where we longitudinally followed them over 300 days. 390 samples were collected at five different time points following their 3 doses of vaccination (**Figure 1A**). The five time points are pre-vaccination baseline (T1), 1 week post dose 1 (T2), 2 weeks post dose 2 (T3), 6-8 months post dose 2 (T4), and 2 weeks post a boost dose 3 (T5). This study design allowed us to monitor the immunological alterations, especially the induction, maintenance, waning and boosting of antigen-specific immune responses to the vaccine in a relatively longer period.

**Figure 1.**
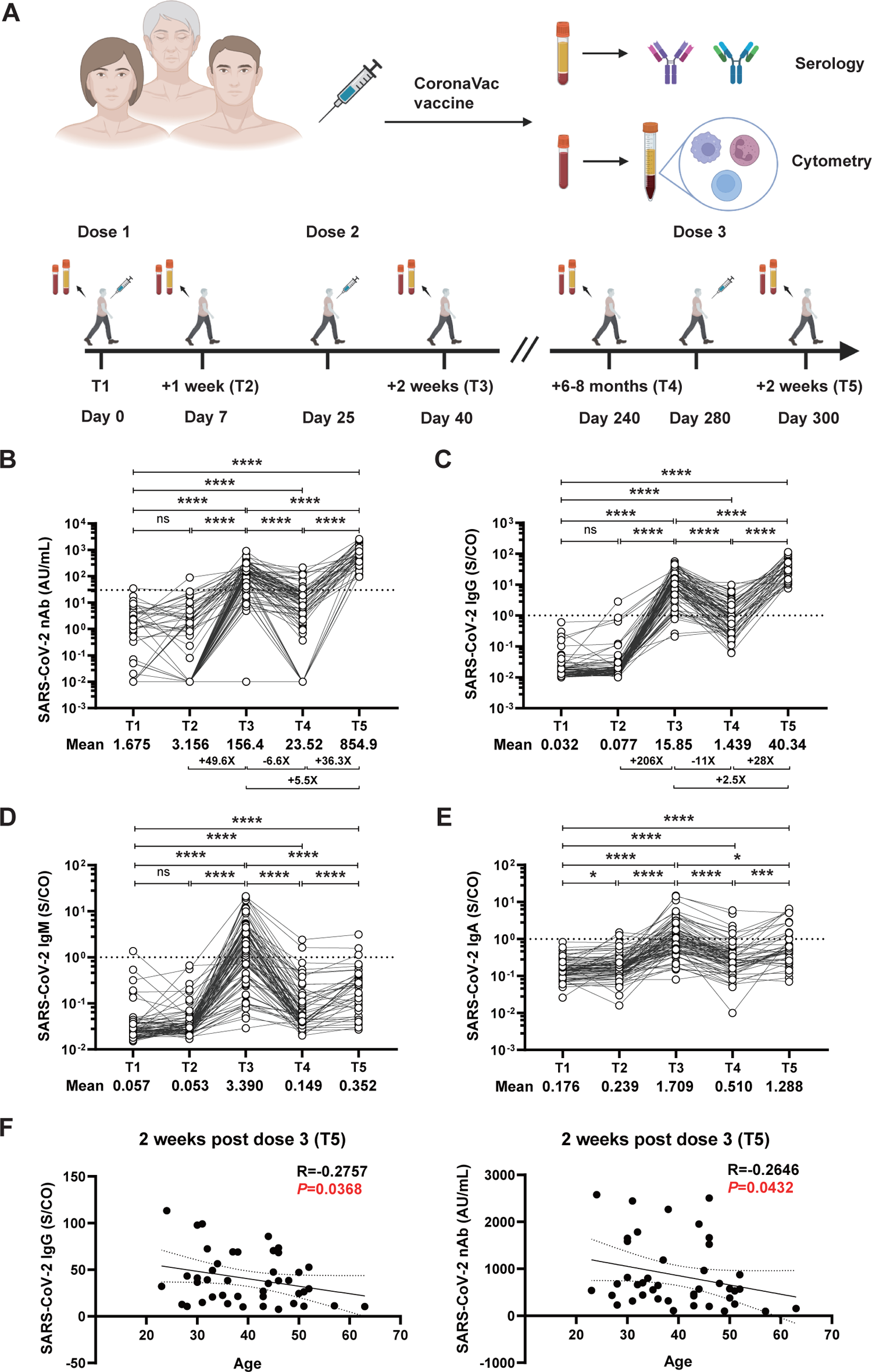
Dynamics of SARS-CoV-2 specific antibody responses. (A) Schematics (Created with BioRender). (B) SARS-CoV-2 nAb titer; (C) Anti-spike IgG titer; (D) Anti-spike IgM; (E) Anti-spike IgA antibodies in vaccinated individuals at five different time points, including pre-vaccination (T1), 1 week after the 1^st^ dose (T2), 2 weeks after the 2^nd^ dose (T3), 6-8 months after the 2^nd^ dose (T4), as well as 2 weeks post the 3^rd^ dose(T5). (F) Correlation between age and SARS-CoV-2 IgG, nAb titers at 2 weeks after the 3^rd^ dose (T5). Each dot represents an individual subject. ‘+X’ indicates fold changes for selected comparisons. ‘–X’ indicates decreased fold changes for selected comparisons. The dashed line indicates the cut-off value, and the samples above the dashed line are considered reactive while below are considered nonreactive. Statistics were calculated using Wilcoxon matched-pairs signed-ranks test for comparison between time points (B, C, D, E). \**P* < 0.05; \*\**P* < 0.01; *** *P* < 0.001; **** *P* < 0.0001; ns, not significant. The two-tailed, non-parametric Spearman’s rank correlation was used (F). *P* and R values were indicated.

At the baseline (T1), all participants had undetectable levels of neutralizing antibodies (nAbs). Consistent with previous reports (10–12), the 2^nd^ dose of CoronaVac vaccine significantly enhanced nAb responses against the wild-type (WT) SARS-CoV-2, with the mean value of nAb rising from only 3.156 AU/mL after the primary dose (T2) to 156.4 AU/mL after the second dose (T3). Different from mRNA vaccine-induced antibody response that remain in a relatively high level 6 months post the 2^nd^ dose of vaccine (13–15), the average nAb antibody-elicited by CoronaVac vaccine dropped rapidly to 23.52 AU/mL after 6-8 months (T4). Importantly, a 3^rd^ dose of CoronaVac vaccine significantly boosted nAb responses, and the seropositivity rate of nAb rapidly reached 100% (**Table 1**). Moreover, a 3^rd^ dose of vaccine also significantly enhanced the magnitude of nAb levels (*P* < 0.0001). The average level of nAb reached 854.9 AU/mL at 2 weeks post dose 3 (T5), a 5.5-fold higher than that at T3 and 36.3-fold higher than that at T4 (**Figure 1B**). Similarly, the seropositivity rate and the amount of anti-Spike IgG increased significantly after the 2^nd^ vaccine dose, with a continued increase post the 3^rd^ vaccination. The mean value of anti-Spike IgG at T5 increased 2.5-fold and 28-fold compared with those at T3 and T4, respectively (**Figure 1C**). Spike-specific IgM (**Figure 1D**) and IgA (**Figure 1E**) displayed a similar kinetics as IgG and nAb during the primary 2-dose vaccine series. By contrast, a 3^rd^ vaccination did not markedly increase IgM and IgA response. The seropositivity rates for anti-Spike IgM and anti-Spike IgA after the 3^rd^ dose (T5) were much lower than that observed after the 2^nd^ dose (T3) (**Table 1**). We also compared the effects of vaccination interval during the primary and secondary dose and found minor impacts on the subsequent immune response **(Figure S1A and S1B)**. Notably, we observed that anti-WT SARS-CoV-2 IgG and nAb after the 3^rd^ dose of vaccination were negatively correlated with age (Figure 1F). Together, these results suggest that the 2^nd^ and 3^rd^ CoronaVac vaccine induced robust nAbs with a 3^rd^ booster further increased the response.

**Table 1.** Spike-specific antibodies detected in CoronaVac vaccinees

|  | T1 | T2 | T3 | T4 | T5 |
| --- | --- | --- | --- | --- | --- |
| No. | 88 | 88 | 88 | 83 | 43 |
| Anti-spike IgG | 0 | 1.136%<br>(1/88) | 95.455%<br>(84/88) | 38.554%<br>(32/83) | 100% |
| Anti-spike IgM | 1.136%<br>(1/88) | 0 | 60.227%<br>(53/88) | 2.410%<br>(2/83) | 6.977%<br>(3/43) |
| Anti-spike IgA | 0 | 2.273%<br>(2/88) | 44.318%<br>(39/88) | 8.434%<br>(7/83) | 27.907%<br>(12/43) |
| Neutralizing antibodies | 1.136%<br>(1/88) | 1.136%<br>(1/88) | 82.955%<br>(73/88) | 15.663%<br>(13/83) | 100% |

The first row indicated the total number of specimens collected at pre-vaccination baseline (T1), 1 week post dose 1 (T2), 2 weeks post dose 2 (T3), 6-8 months post dose 2 (T4), and 2 weeks post a boost dose 3 (T5). The remaining parts showed the positive rates of serum anti-spike IgG, anti-spike IgM, anti-spike IgA and neutralizing antibodies at five time points, and the brackets indicated the specific number of antibody positive at each time point.

### Plasma neutralization against variants of concern

To examine the neutralizing efficacy and breadth of antibodies elicited by CoronaVac vaccine, especially their evolution and persistence following each vaccination, pseudovirus neutralization was applied and 50% inhibitory dose (ID_50_) was used to calculate the plasma neutralization against ancestral SARS-CoV-2 (WT), Omicron B.1.1.529 and BA.2 variants using 72 longitudinal samples from 6 randomly selected individuals. All samples tested showed no neutralization activities (ID_50_=15) against all three strains following 1^st^ dose of CoronaVac vaccine (**Figure 2A**). While this neutralization activities were significantly improved following the 2^nd^ dose of vaccination as most of the individuals (5/6) showed significantly increased ID_50_, ranging from 35 to 128, against ancestral SARS-CoV-2 (**Figure 2B**). Nearly all individuals showed poor plasma neutralizations against Omicron B.1.1.529 and BA.2 variant following 2^nd^ vaccine dose (**Figure 2B**). Although half of the individuals (3/6) maintained their plasma neutralizing activities against ancestral (WT) SARS-CoV-2 strain, the overall ID_50_ waned sharply 6 months post 2^nd^ dose of CoronaVac vaccine (ID_50_ ranging from 15-38) (**Figure 2C**). Interestingly, after a 3^rd^ boost of CoronaVac vaccine, both the efficacy and breadth of plasma neutralization were markedly improved (**Figure 2D**). Importantly, 4 out 6 individuals developed adequate level of cross-neutralizing activities against Omicron sub-lineage B.1.1.529 and BA.2 (**Figure 2D**), which was not seen in individuals prior to receive their 3^rd^ dose of CoronaVac vaccine. In line with the evidence from mRNA vaccine induced neutralizing antibody protection, a 3^rd^ booster of CoronaVac vaccine was necessary to significantly increase the efficacy and breadth of the protective neutralizing antibody response (16).

**Figure 2.**
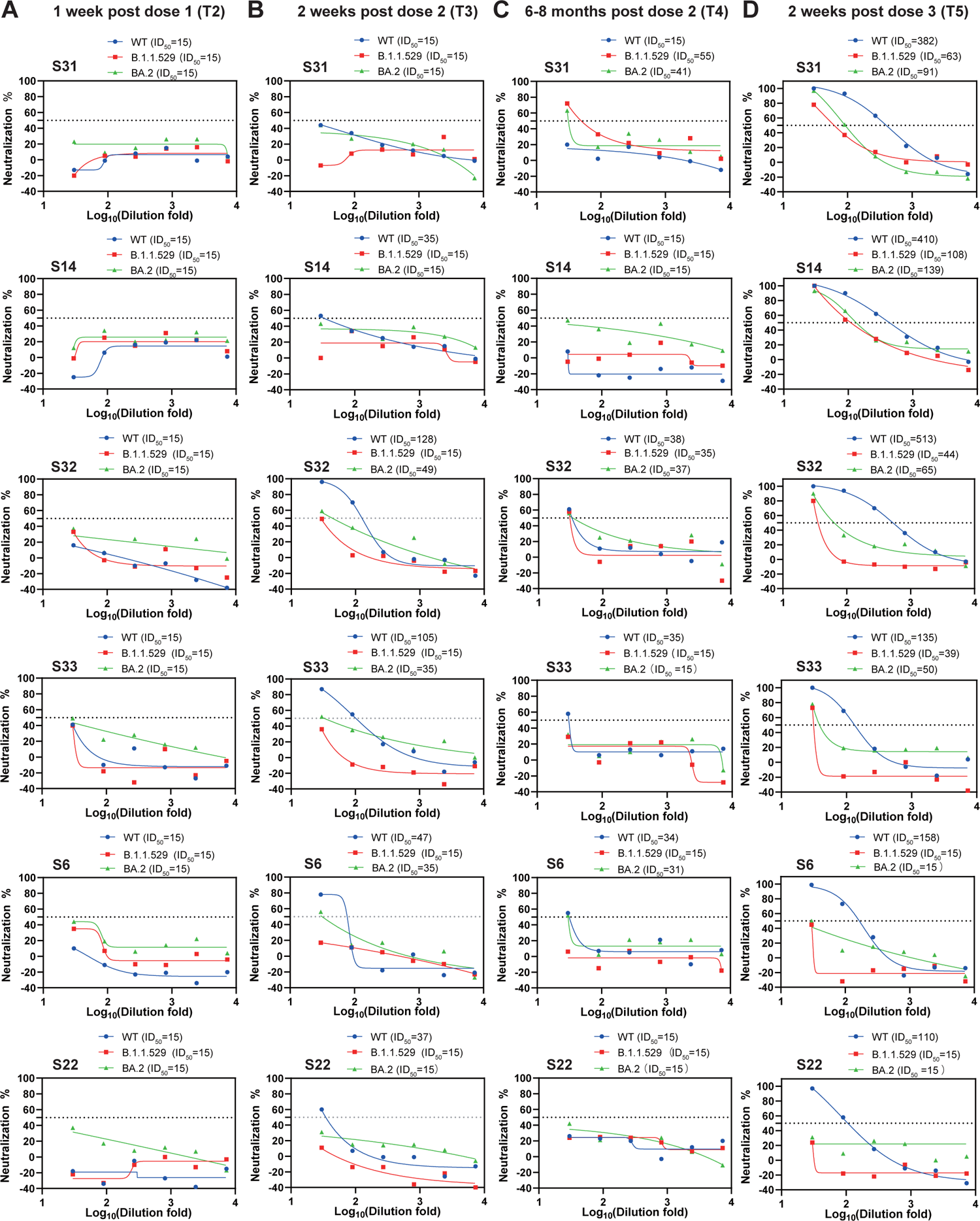
Neutralizing activities against SARS-CoV-2 WT and VOC. Pseudovirus neutralization titers against SARS-CoV-2 WT, B.1.1.529, and BA.2 variant using plasma samples from six randomly selected individuals. The sera were collected at 1 week after the 1^st^ dose (T2) (A), 2 weeks after the 2^nd^ dose (T3) (B), 6-8 months after the 2^nd^ dose (T4) (C), and 2 weeks after the 3^rd^ dose (T5) (D), respectively. The horizontal dash lines indicate 50% of the pseudovirus neutralization (ID_50_).**^10^**

Little is known about the neutralization potential of emerging Omicron sub-variants such as BA.4/BA.5 and BA.2.75.2 by inactivated vaccines. We set out to address this urgent question by testing samples from 10 individuals received the 3^rd^ vaccine booster. 4 out of 10 donors showed broad neutralization against all four Omicron sub-variants (**Figure. 3A**). 6 out of 10 donors exhibited robust neutralizing antibody response against BA.4/BA.5 (**Figure 3A**). Interestingly, plasmas from 8 out 10 donors neutralized the BA.2.75.2 (**Figure 3A**). ID_50_ of 3^rd^booster-elicited antibodies against BA.4/BA.5 and BA.2.75.2 is lower than that of against B.1.1.529 and BA.2 (**Figure 3B**). However, such reduction is relatively minor comparing to the ID_50_ against the ancestral SARS-CoV-2 (**Figure 3B**), which is consistent with the most recent reports (17, 18), BA.2.75.2 is suggested to be more immune evasive than BA.5 (19), our results imply that the 3^rd^ booster of CoronaVac vaccine generates modest but relatively broad antibody protection against the Omicron sub-variants that is currently circulating.

**Figure 3.**
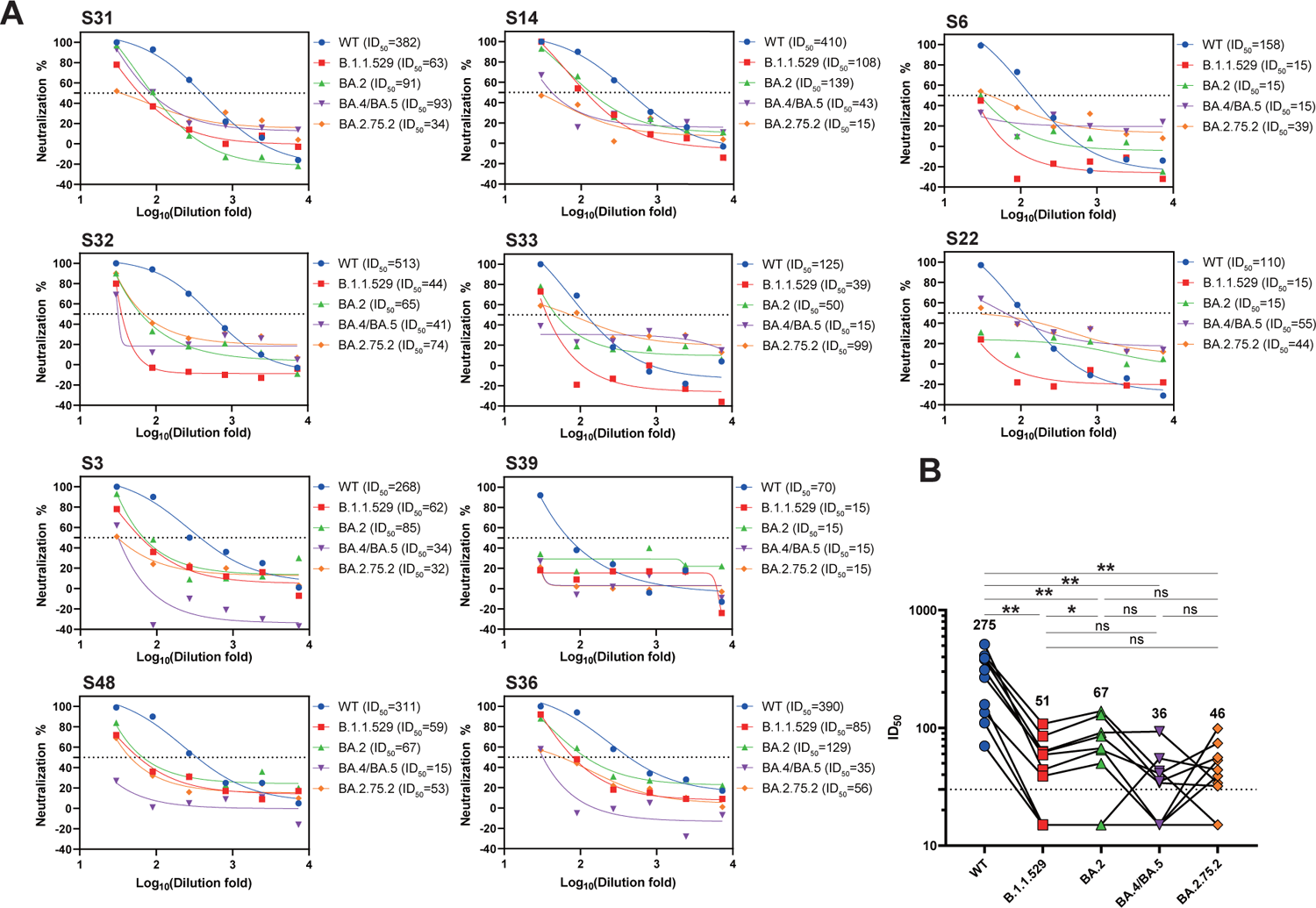
Protection of neutralizing antibody against SARS-CoV-2 variants after the 3^rd^ dose of CoronaVac. (A) Neutralizing activities against SARS-CoV-2 WT, B.1.1.529, BA.2, BA.4/BA.5 and BA.2.75.2 variants of plasma samples from 10 randomly selected individuals who received the 3^rd^ dose of CoronaVac vaccination. The horizontal dash lines indicate 50% of the pseudovirus neutralization (ID_50_). Patient numbers and ID_50_ of different variants are shown at the top of each graph. (B) Neutralizing antibody titers (indicated as ID_50_) against SARS-CoV-2 WT, B.1.1.529, BA.2, BA.4/BA.5 and BA.2.75.2 variants measured post the 3^rd^ vaccination. Statistical significance was determined using Wilcoxon matched-pairs signed-ranks test for comparison between SARS-CoV-2 variants. *, *P* < 0.05; **, *P* < 0.01; ns, not significant.

### Polyclonal circulating CD4^+^ T cell response

Next, we set out to evaluate polyclonal peripheral CD4^+^ T cells in our longitudinal cohorts and determine how CD4^+^ T cell subsets might evolve by 3 doses of CoronaVac vaccine. Multicolor flow cytometry was used to measure the frequency of different CD4^+^ T cell subsets. The specific gating strategies based on the combination of signature surface molecules for lymphocytes were shown in **(Figure S2A)**. Polyclonal memory and naïve CD4^+^ T cell were identified based on CD45RA and CCR7 (**Figure 4A**) (20). Compared to the cells before vaccination (T1), we observed a marked increase of effector memory (EM) CD4^+^ T cells shortly after each dose of vaccination (T2, T3, T5), accompanied by declined frequencies of central memory (CM) CD4^+^ T cells (**Figures 4B and 4C**). While naïve CD4^+^ T cell frequency was rather stable over the course of our longitudinal follow-up (**Figures 4B and 4C**). When evaluating the functional CD4^+^ T cell subsets, we found relatively minor but significant increase of T_H_1 and decrease of T_H_2 cells after the 2^nd^ dose (T3) compared with baseline (T1), while negligible changes in T_H_17 cells **(Figure S2B).** T follicular helper (T_FH_) cells provide help to B cells to promote germinal center (GC) selection of memory and plasma B cells in health and disease (21–23). Post 1^st^ dose of CoronaVac, we observed comparable circulating T_FH_ (cT_FH_) cells in vaccinees, while this frequency of polyclonal cT_FH_ cells were significantly increased by a 2^nd^ dose of vaccine (**Figures 4D**). Polyclonal cT_FH_ cell level was declined over the following 6-8 months and maintained at similar level upon a 3^rd^ boost of CoronaVac vaccine (**Figure 4D**). Interestingly, when measuring cT_FH_ cell subsets featured by the differential expression of CXCR3 and CCR6 (24), We observed a significant change in the composition of these subsets following administration of a 2^nd^ and 3^rd^ dose of the vaccine (**Figure 4E**). With that being observed, the proportion of CXCR3-expressing cT_FH_1 cells were particularly increased by a 2^nd^ dose and 3^rd^ dose of vaccine, while the proportion of cT_FH_2 cells were largely reduced among cT_FH_ cells (**Figure 4F**). Moreover, cT_FH_17 cells shared similar trends with that of found in cT_FH_1 cells (**Figure 4F**). The total frequencies of each cT_FH_ subsets were also evaluated and consistently, we observed increased cT_FH_1 cells by the 2^nd^ and 3^rd^ vaccination comparing to baseline, while comparable or subtle differences with regards to the cT_FH_2 and cT_FH_17 cells (**Figure 4G**).

**Figure 4.**
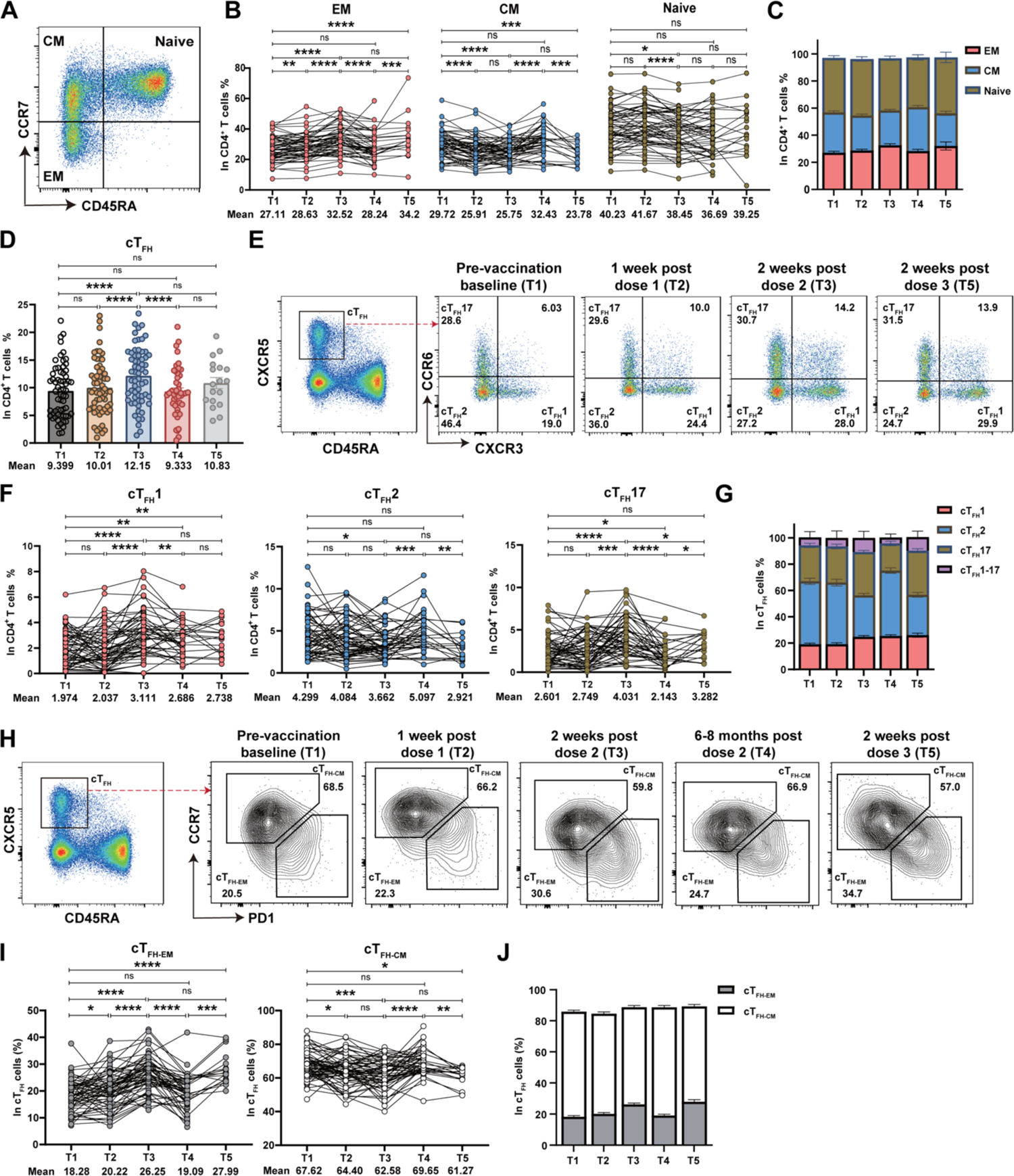
Characterization of polyclonal peripheral CD4^+^ T cells. PBMCs collected from vaccinated donors (n = 63) at five different time points (T1-T5) were analyzed by 24-color flow cytometry. (A) Representative FACS plots of CD4^+^ T cell memory subsets and CD4^+^ naïve T cells defined by CD45RA and CCR7. (B) Statistical analysis of the frequency of polyclonal effector memory (EM), central memory (CM) and naïve CD4^+^ T cells at five time points. (C) Composition of polyclonal EM, CM and naïve CD4^+^ T cells from vaccinated individuals at five time points. Data are the same as in (B). (D) Statistical analysis of the frequency of polyclonal cT_FH_ cells at five time points. (E) Representative FACS disgrams of cT_FH_ subsets grouped by CCR6 and CXCR3 at T1, T2, T3 and T5. (F) Longitudinal frequencies of polyclonal cT_FH_1, cT_FH_2 and cT_FH_17 cells measured by flow cytometry at five time points. (G) The proportion of cT_FH_1, cT_FH_2, cT_FH_17 and cT_FH_1-17 cells in polyclonal cT_FH_ cells at five time points. Data are the same as in (F). (H) Representative FACS maps of cT_FH-EM_ and cT_FH-CM_ cell subsets gated by PD1 and CCR7 in cT_FH_ cells at five time points. (I) Statistical analysis showing the differences of the frequencies of CCR7^hi^PD1^-^ cT_FH-CM_ and CCR7^lo^PD1^+^ cT_FH-EM_ cells at five time points. (J) The proportion of cT_FH-EM_ and cT_FH-CM_ cells in polyclonal cT_FH_ cells at five time points. Data are the same as in (I). Each dot represents an individual subject. Bars represent the mean values with SEM. Statistics were calculated using Wilcoxon matched-pairs signed-ranks test for comparison between time points (B, D, F, I). *, *P* < 0.05; **, *P* < 0.01; ***, *P* < 0.001; ****, *P* < 0.0001; ns, not significant.

Next, we examined the effector and memory cT_FH_ cell response following each vaccine dose, by looking into the surface expression of CCR7 and PD-1 on cT_FH_ cells (24, 25). The frequency of CCR7^lo^PD1^hi^ effector-memory circulating T_FH_ (cT_FH-EM_) cells were significantly increased by the 2^nd^ and 3^rd^ dose of vaccine (**Figure 4H and 4I**). While the frequency of cT_FH-EM_ cells dropped markedly 6-8 months after the 2^nd^ dose of vaccination (**Figure 4I and 4J**). Correspondingly, the frequency of CCR7^hi^PD1^lo^ central-memory cT_FH_ (cT_FH-CM_) cells increased significantly over the time course of 6-8 months post the 2^nd^ vaccination (**Figure 4I and 4J**). The kinetics and the rapid alterations of effector and memory cT_FH_ cells following a vaccine administration further support circulating T_FH_ cell as the key biomarker when evaluating the effectiveness and longevity of a vaccine response.

### T_FH_1 cells represent the effector T_FH_ cells that effectively respond to the vaccination

Both cT_FH_1 and cT_FH_17 cells have been shown to correlate with antibody responses induced by SARS-CoV-2 infection or vaccination (26). However, how these T_FH_ cell subsets evolve and persist remains largely unknown. Human T_H_17 cells phenotypically resembled memory T cell in autoimmunity and anti-tumor response and showed higher capacity for proliferative self-renewal and plasticity to interconvert into other CD4^+^ T cell subsets (27). By contrast, T_H_1 cells are more terminally differentiated during viral infection (28). To understand whether cT_FH_1 cells and cT_FH_17 cells possess different effector and memory potential, we performed Pearson correlation coefficient analysis between subsets of cT_FH_ cells and effector/memory cT_FH_ cells in donors after their 2^nd^ vaccination and 3^rd^ vaccine booster (**Figure 5A**). Strikingly, we found strong positive correlations between cT_FH_1 cells and cT_FH-EM_ cells after both vaccinations. Similar positive correlations were noticed between cT_FH_17 cells and cT_FH-CM_ cells (**Figure 5A**). In contrast, cT_FH_1 cells showed significant negative association with cT_FH-CM_ cells, with the same trend between cT_FH_17 cells and cT_FH-EM_ cells. To further confirm the dominance of cT_FH-EM_ cells in cT_FH_1 cells while cT_FH-CM_ cells in cT_FH_17 cells, we evaluated the frequency of EM and CM cells in each cT_FH_ subsets at five different time points in 63 vaccinees (**Figure 5B**). We found that although more cT_FH_1 cells were CM cells before vaccination, markedly increased proportion of EM cells were observed in cT_FH_1 cells following the 2^nd^ and 3^rd^ vaccination (30% to 39.61%, p<0.0001; 30% to 40.24%, p=0.0004) (**Figure 5C**). These proportion of EM cells sharply declined 6-8 months after the 2^nd^ vaccination (39.61 to 32.41%, p<0.0001), suggesting a short-lived phenotype of cT_FH_1-EM cells. Nevertheless, a 3^rd^ booster of vaccine rapidly reinvigorated the frequency of cT_FH_1-EM cells to 40.24%. In contrast, over 70% of cT_FH_17 cells were CM cells, with as little as 10-20% cT_FH_17 cells were EM cells over the course of 3 vaccine administrations (**Figure 5C**). Notably, the proportion of CM cells in cT_FH_17 cells remained intact over the time course of 3 vaccine administrations, suggesting a relatively long-lived phenotype of cT_FH_17 cells (**Figure 5C**). Taken together, our data suggest that T_FH_1 cells likely constitute the majority of effector T_FH_ cells that effectively respond to sequential vaccination, but are short-lived. Conversely, T_FH_17 cells may resemble the most long-lived memory T_FH_ cells induced by the vaccine.

**Figure 5.**
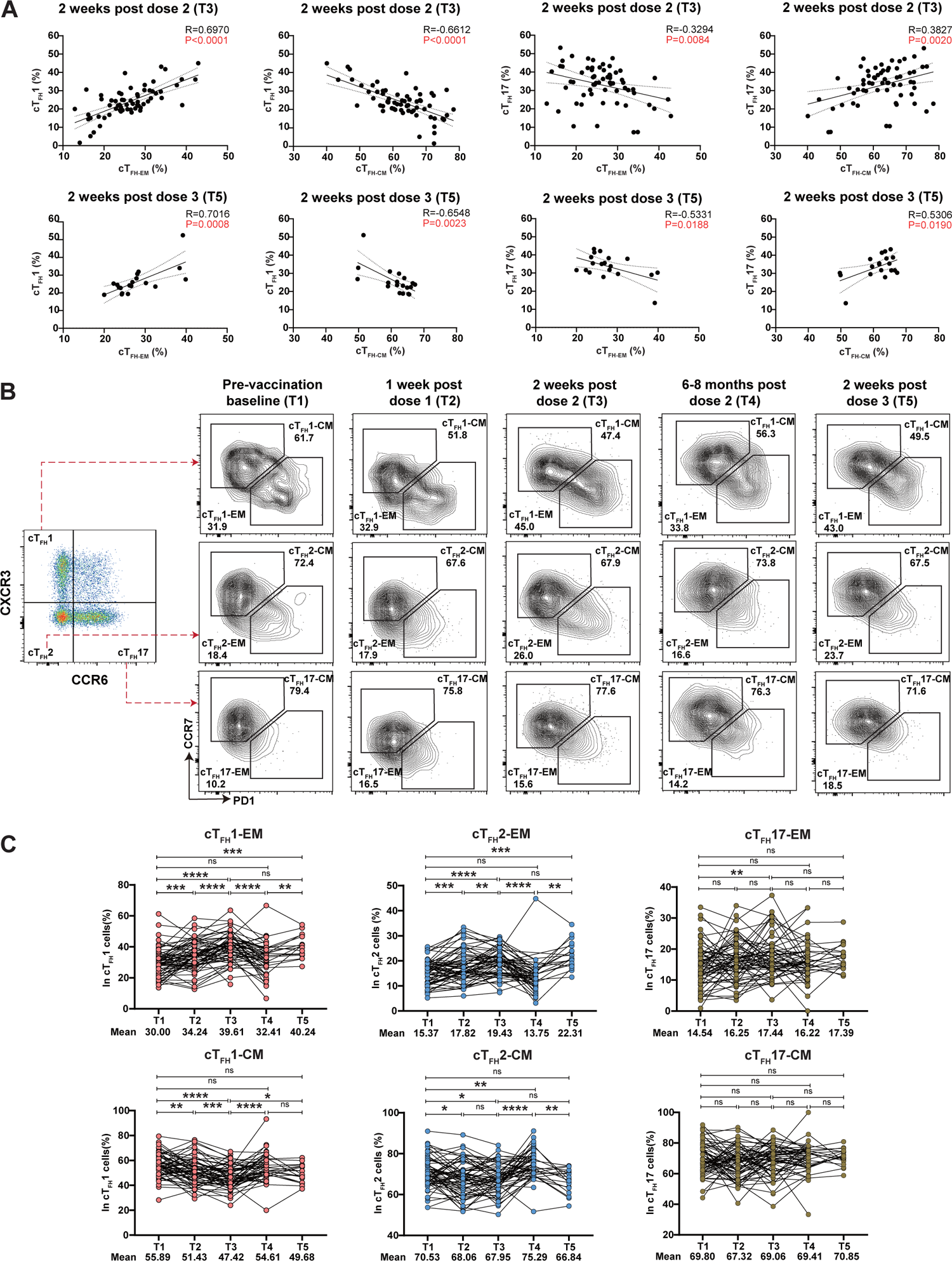
Characterization of effector and memory cT_FH_ cells following vaccination. (A) Correlation analysis between cT_FH_1, cT_FH_17 and cT_FH-EM_, cT_FH-CM_ cell frequencies at 2 weeks post 2 (T3) and 2 weeks post dose 3 (T5). (B) FACS plots showing the representative cT_FH_1-EM, cT_FH_1-CM, cT_FH_2-EM, cT_FH_2-CM, cT_FH_17-EM and cT_FH_17-CM cells gating from cT_FH_1, cT_FH_2, cT_FH_17 cells by CCR7 and PD1 at five time points. (C) Frequencies of cT_FH_1-EM, cT_FH_1-CM, cT_FH_2-EM, cT_FH_2-CM, cT_FH_17-EM and cT_FH_17-CM cells within cT_FH_1, cT_FH_2 and cT_FH_17 cells at five time points. Each dot represents an individual subject. The 2-tailed Pearson correlation test was used (A). *P* and R values were indicated (A). Statistics were calculated using Wilcoxon matched-pairs signed-ranks test for comparison between time points (C). *, *P* < 0.05; **, *P* < 0.01; ***, *P* < 0.001; ****, *P* < 0.0001; ns, not significant (C).

### Spike-specific CD4^+^ T cell response

To determine whether CoronaVac inactivated vaccine can induce durable antigen-specific memory CD4^+^ T cell responses, we utilized activation-induced marker (AIM) assay and evaluated the SARS-CoV-2 spike-specific response. PBMCs were stimulated with SARS-CoV-2 spike peptides, containing a pool of both S1 and S2 peptides, or SEB as positive control. AIM^+^ CD4^+^ T cells were defined by dual expression of CD25 and HLA-DR (**Figure 6A**) (29–31). For spike-specific CD4^+^ T cell response from each time point, PBMCs were also treated with DMSO to define the spike-positive population (**Figure 6A**). Full gating strategies are provided in **(Figure S3)**. The frequency of AIM^+^ CD4^+^ T cells increased slightly post 1^st^ dose of vaccine (T2), with around 2-folds further increase post 2^nd^ dose of vaccine (T3) comparing to that of in baseline (T1) (**Figures 6A and 6B**). These results indicate robust induction of SARS-CoV-2 spike-specific CD4^+^ T cell responses after vaccination. Moreover, 6-8 months after the 2^nd^ vaccine dose (T4), the frequency of AIM^+^ CD4^+^ T cells was shapely deceased for about 1.4 folds comparing to that of 2 weeks after the 2^nd^ vaccination (T3). Notwithstanding the AIM^+^ CD4^+^ T cells still maintained at a detectable level and higher than that of in the baseline (**Figure 6B**). As expected, a 3^rd^ booster of CoronaVac reinvigorated the AIM^+^ CD4^+^ T cells to a comparable level to that of soon after the 2^nd^ vaccination (T3) (**Figure 6B**). The proportion of EM cells in the total spike-specific CD4^+^ T cells was maintained at a high level at T2, T3, T4 and T5, while the proportion of CM cells increased significantly at 6-8 months after the second dose (T4) (**Figure 6C**).

**Figure 6.**
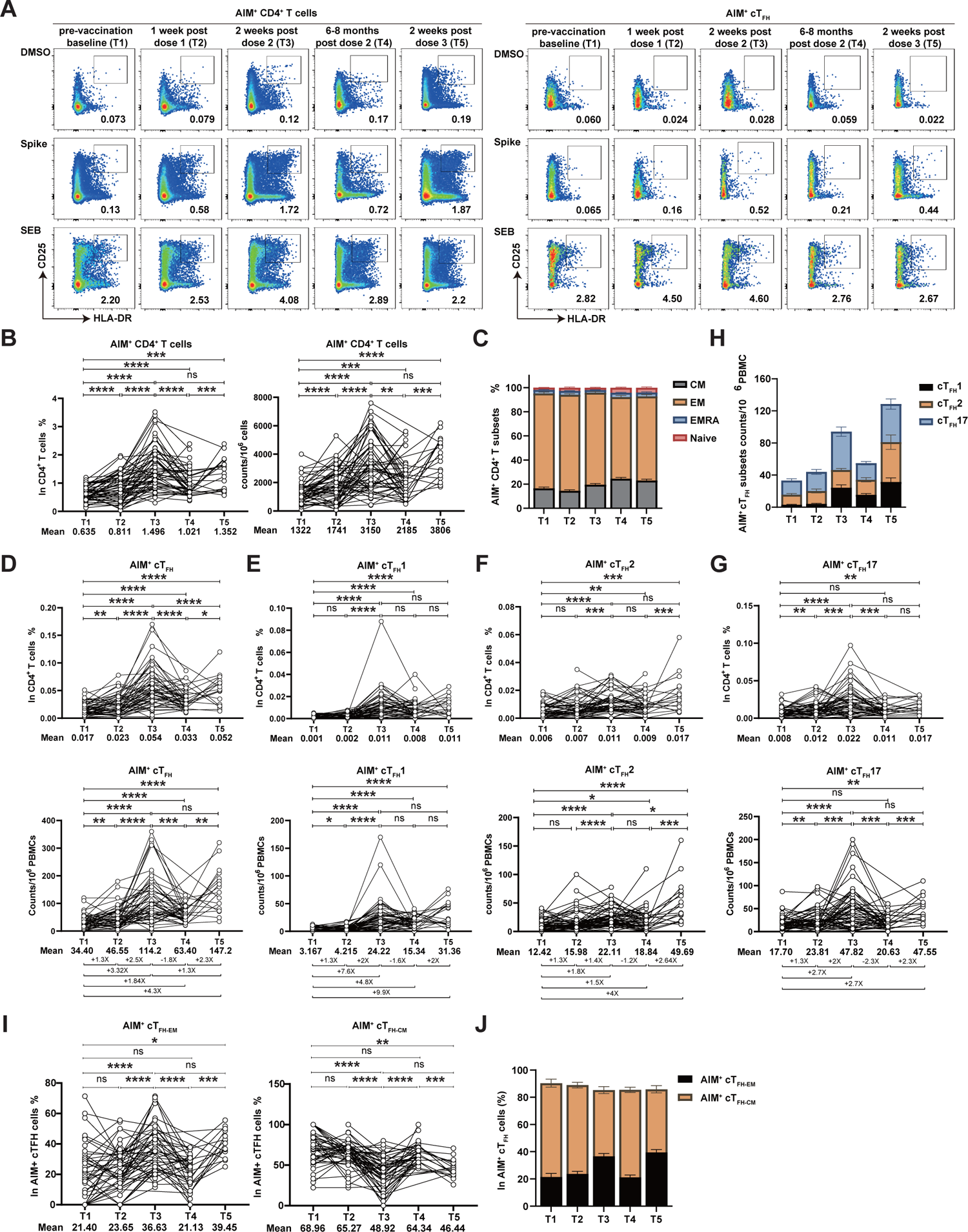
CoronaVac-induced spike-specific memory T follicular helper cells. PBMCs collected from vaccinated donors (n = 63) at five different time points (T1-T5) were *ex vivo* stimulated with SARS-CoV-2 spike protein (S1+S2, 2 ug/mL, SinoBiological) in 5% CO_2_ at 37°C for 24h. SEB (1 ug/mL, Toxin Technology) were used as positive control. (A) Representative FACS plots of AIM^+^ CD4^+^ T (HLA-DR^+^CD25^+^) cells and AIM^+^ cT_FH_ (CXCR5^+^ HLA-DR^+^CD25^+^) cells at five time points. The frequencies of AIM^+^ CD4^+^ T cells (B), AIM^+^ cT_FH_ cells (D), AIM^+^ cT_FH_1 cells (E), AIM^+^ cT_FH_2 cells (F) and AIM^+^ T_FH_17 cells (G) were shown by the percentage in total CD4^+^ T cells and cell numbers in 10^6^ PBMCs. (C) The frequencies of AIM^+^ EM, CM, EMRA and naïve CD4^+^ T cells in AIM^+^ CD4^+^ T cells at five time points. (H) The cell numbers of AIM^+^ cT_FH_1, cT_FH_2 and cT_FH_17 cells at five time points. Data are the same as in (E, F and G). (I) Statistical analysis showing the alteration of the frequencies of AIM^+^ cT_FH-EM_ and AIM^+^ cT_FH-CM_ cells at five time points. (J) The proportion of AIM^+^ cT_FH-EM_ and AIM^+^ cT_FH-CM_ cells in AIM^+^ cT_FH_ cells at five time points. Data are the same as in (I). Each dot represents an individual subject. Bars represent the mean values with SEM. Statistics were calculated using Wilcoxon matched-pairs signed-ranks test for comparison between time points (B, D, E, F, G) *, *P* < 0.05; **, *P* < 0.01; ***, *P* < 0.001; ****, *P* < 0.0001; ns, not significant.

To further assess the functionality of CoronaVac-induced CD4^+^ T cell responses, we characterized the SARS-CoV-2 spike-specific circulating T_FH_ cells (CXCR5^+^HLA-DR^+^CD25^+^CD4^+^) (**Figure 6A, Right)**. Notably, although the virus-specific IgG and nAbs were low to non-detectable 1 week post 1^st^ vaccination (T2), we found slightly increased frequency of spike-specific cT_FH_ cells relative to the baseline (p=0.0043). Similar to the total spike-specific CD4^+^ T cells, the frequency and number of spike-specific cT_FH_ cells were further increased post 2^nd^ dose of vaccine but steeply decreased 6-8 months later (**Figure 6D**). The 3^rd^ dose of CoronaVac vaccination enhanced the spike-specific cT_FH_ response to the similar magnitude shortly after a 2^nd^ vaccine booster (**Figure 6D**). Spike-specific cT_FH_ cell were further examined to understand the memory potential and superiority of its functional subsets, which might support the humoral immune response differently. Interestingly, we found that the magnitude of spike-specific cT_FH_1 response were relatively constant after boosted by a dose of CoronaVac vaccine, with no significant decrease or increase of cT_FH_1 cells over 6-8 months and by a 3^rd^ booster (**Figure 6E and 6H**). Interestingly, highest spike-specific cT_FH_2 cell numbers were found after a 3^rd^ boost of vaccine (**Figure 6F and 6H**), which were not seen in polyclonal cT_FH_ cells (**Figure 4F**). Spike-specific cT_FH_17 cells, however, share a similar kinetics to polyclonal cT_FH_17 cells, where the 2^nd^ dose of vaccine significantly mounted spike-specific cT_FH_17 cell response which was declined over 6-8 months but reinvigorated by a 3^rd^ vaccine booster (**Figure 6G and 6H**). Of note, when evaluating the memory potential of spike-specific cT_FH_ cells by CCR7 and PD1, we found significantly increased frequency of spike-specific CCR7^lo^PD1^hi^ cT_FH-EM_ cells by the 2^nd^ dose (**Figure 6I**). Similar to the polyclonal T_FH_ cell response in (**Figure 4I**), the frequency of spike-specific cT_FH-EM_ cells waned 6-8 months after the 2^nd^ dose of vaccination but reinvigorated by the 3^rd^ dose of vaccine booster (**Figure 6I**). Correspondingly, spike-specific cT_FH-CM_ cells were markedly reduced by the 2^nd^or the 3^rd^ vaccination (**Figure 6I and 6J**). Together, these data suggest that the 2^nd^ dose and 3^rd^ dose of CoronaVac vaccine induce robust spike-specific CD4^+^ T cell and cT_FH_ cell responses, with distinctive patterns shared by different functional subsets.

In line with the spike-specific CD4^+^ T cell response, markedly increased spike-specific IL-2 and IFN-γ producing CD4^+^ T cells were noticed 7 days after CoronaVac vaccination (**Figure S4A & Figure 7A**). The 2^nd^ and 3^rd^ CoronaVac booster further increased the frequencies of IL-2^+^ and IFN-γ^+^ CD4^+^ T cells relative to baseline and the 1^st^ immunization (**Figure S4B-C, Figure 7A**). We further assessed the cytokine production from spike-specific T_H_1 cells by gating on the CXCR3^+^ CD4^+^ T cells and found similar trends of IL-2 and IFN-γ production promoted by multiple CoronaVac vaccinations (**Figure 7B**). These data, together, indicates that the 2^nd^ and 3^rd^ homologues CoronaVac booster elicit functional spike-specific CD4^+^ T cells necessary for regulating anti-viral responses.

**Figure 7.**
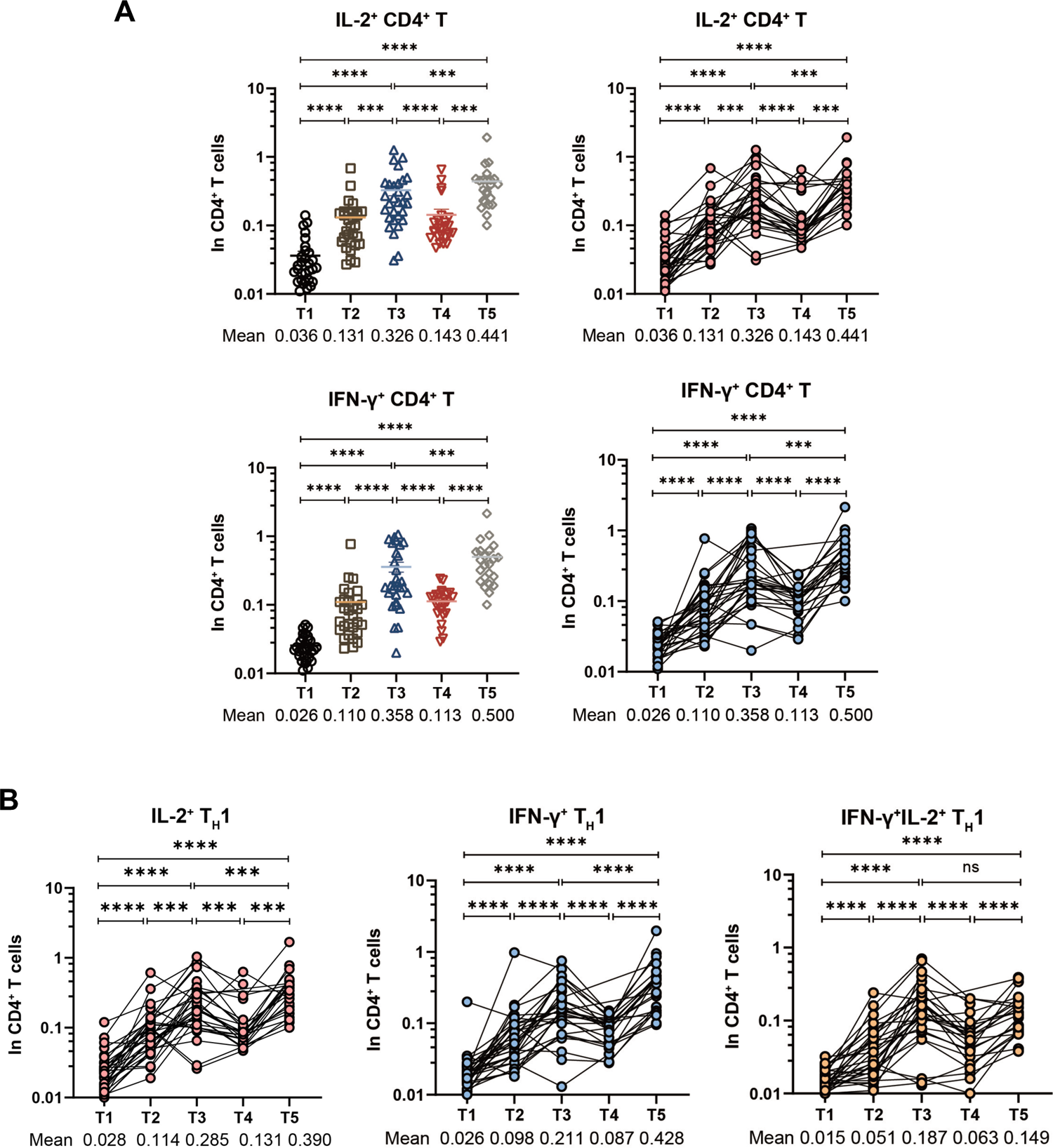
Cytokine-producing spike-specific CD4+ T cell responses. PBMCs collected from vaccinated donors at five different time points (T1-T3, n=30; T4, n=25; T5, n=23) were *ex vivo* stimulated with SARS-CoV-2 spike protein (S1+S2, 2 ug/mL, SinoBiological) in 5% CO_2_ at 37°C for 24h. (A) Frequencies of IL-2^+^ (up), IFN-γ^+^ (bottom) spike-specific CD4^+^ T cells detected after SARS-CoV-2 peptide stimulation at different timepoints. (B) Frequencies of IL-2^+^ (up), IFN-γ^+^ (bottom) spike-specific T_H_1 cells observed following SARS-CoV-2 peptide stimulation at different timepoints. Each dot represents an individual subject. Statistics were calculated using Wilcoxon matched-pairs signed-ranks test for comparison between time points (A, B) *, *P* < 0.05; **, *P* < 0.01; ***, *P* < 0.001; ****, *P* < 0.0001; ns, not significant.

### Correlations between CD4^+^ T cells and antibody responses following CoronaVac vaccination

To determine the relationship between CD4^+^ T cells and the production of SARS-CoV-2 antibodies after CoronaVac vaccination, we firstly analyzed the polyclonal circulating CD4^+^ T subsets and SARS-CoV-2 antibodies after the 2^nd^ and 3^rd^ vaccine dose (T3 and T5). The correlation matrix analysis using non-parametric Spearman’s rank test revealed that there was no statistical significance on the correlation between polyclonal cT_FH_ cells and antibody titers after the 2^nd^ dose (T3). Other CD4^+^ T cell subsets also showed negligible associations **(Figure S5A, left)**. By contrast, polyclonal cT_FH_ cells were positively correlated with both SARS-CoV-2 spike-specific IgG (R = 0.4876, *P* = 0.0401) and IgM (R = 0.5145, *P* = 0.0289) after the 3^rd^ dose of CoronaVac vaccine (T5) **(Figure S5A, right and Figure S5B)**. Total cT_FH_ cells and the level of SARS-CoV-2 nAb showed borderline correlations (R = 0.3765, *P* = 0.0618) **(Figure S5B)**. Interestingly, we found that both polyclonal cT_FH_1 (R = 0.4947, *P* = 0.0369) **(Figure S5C)** and cT_FH_17 (R = 0.5679, *P* = 0.0140) **(Figure S5E)** cells were positively correlated with viral-specific IgM after the 3^rd^ dose (T5), while this was not for cT_FH_2 cells **(Figure S5D).**

Next, we evaluated the relationship between spike-specific AIM^+^ CD4^+^ T cell subsets and SARS-CoV-2 antibody titers after the 2^nd^ dose (T3) and 3^rd^ dose of vaccines (T5) (**Figure 8A**). After multiple corrections with both Spearman’s rank correlation coefficient test and Pearson correlation coefficient test, we found no correlation between AIM^+^ CD4^+^ T cell numbers and viral-specific antibody titers at T3 and T5 (**Figure 8A**). Interestingly, there was a positive correlation between spike-specific cT_FH_ numbers and SARS-CoV-2 IgG (R = 0.2863, *P* = 0.0266) and nAb (R = 0.2393, *P* = 0.0656) titers after the 2^nd^ dose (T3) (**Figure 8B**). Similar trends were also found after the 3^rd^ dose (T5), but the results were not statistical different (**Figure 8B**). In addition, we observed that spike-specific CXCR3^+^ cT_FH_1 cell numbers positively associated with spike-specific IgG (*P* = 0.0592) and nAb titers (*P* = 0.0189) at T3 (**Figure 8C**). The trends still held at T5, although statistically this was not significant (**Figure 8C**). By contrast, there were no correlations between the number of spike-specific cT_FH_2 cells and IgG and nAb levels after the 2^nd^ (T3) and 3^rd^ (T5) dose (**Figure 8D**). It is worth mentioning that AIM^+^ cT_FH_17 cell numbers were positively correlated with SARS-CoV-2 IgG (R = 0.2740, *P* = 0.0341) after the 2^nd^ dose (T2), but not after the 3^rd^ dose (**Figure 8E**). Together, our results suggest that the 2^nd^ and 3^rd^ dose of CoronaVac vaccine induced spike-specific cT_FH_ cells closely associate with serum antibody response and are capable of supporting humoral immune response.

**Figure 8.**
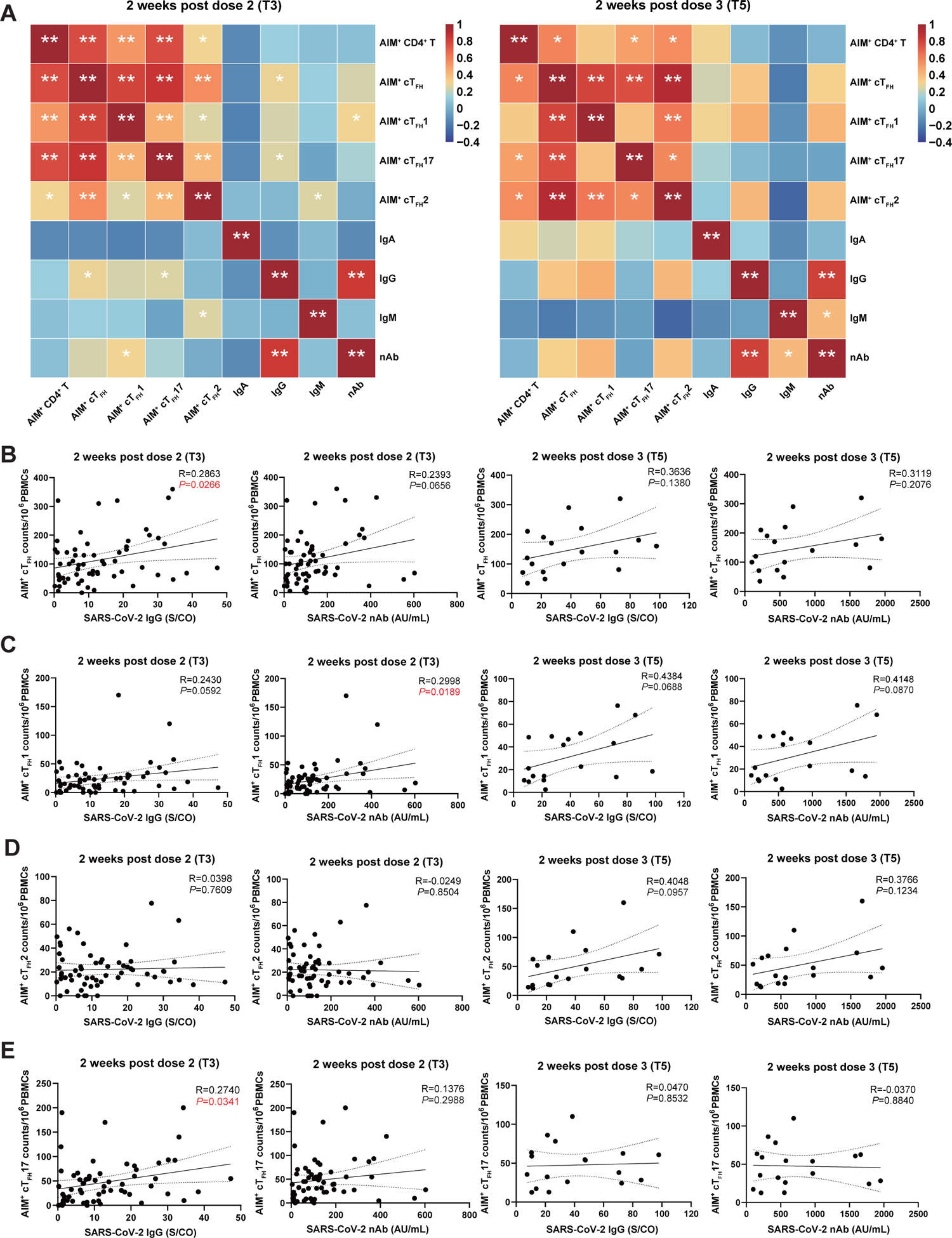
Correlations between CD4^+^ T cells and antibody responses following CoronaVac vaccination. (A) The correlation matrix analysis showed the correlation between AIM^+^ CD4^+^ T cell subsets and SARS-CoV-2 antibodies after the 2^nd^ (T3) and 3^rd^ (T5) dose. Red showed positive correlation, blue represented negative correlation. The color intensity showed the proportion to the correlation coefficients. Correlation analysis between AIM^+^ cT_FH_ (B), AIM^+^ cT_FH_1 (C), AIM^+^ cT_FH_2 (D), AIM^+^ cT_FH_17 (E) cell numbers and SARS-CoV-2-specific IgG, nAb titers at T3 and T5. Each dot represents an individual subject. The two-tailed, non-parametric Pearson and Spearman’s rank correlation test were used, and results corrected post both analyses were showed (A, B, C, D, E). \**P* < 0.05; \*\**P* < 0.01 (A). *P* and R values were indicated (B, C, D, E).

## Discussion

More than 2 billion doses of CoronaVac have been administered in more than 40 countries (32). Recent evidence has shown that a homologous 3^rd^ dose of CoronaVac was associated with further increased SARS-CoV-2-specific antibodies (2–4). Although neutralizing antibody titers were found to be lower in vaccine recipients who received a 3^rd^ booster dose of CoronaVac compared to those who received three doses of mRNA vaccines (33), in general, the 2^nd^ and 3^rd^ doses of CoronaVac were effective in preventing COVID-19-related mortality (74.8% for those aged >65; 80.7% for those aged 50-64; 82.7% for those aged 18-51) and severe complications (58.9% for those aged >65; 67.1% for those aged 50-64; 77.8% for those aged 18-51) (34). Consistent reductions in risk were observed with a 3^rd^ booster dose of CoronaVac (33–36). However, it remains unclear how a 3^rd^ CoronaVac vaccine dose affects the magnitude and quality of immune responses, particularly against the highly divergent variant Omicron. Here, we longitudinally evaluated the CoronaVac vaccine elicited antibody and CD4^+^ T cell responses for 300 days. This allowed us to fill in the knowledge gap of whether the inactivated SARS-CoV-2 vaccine may induce persistent and high quality humoral immune response against Omicron subvariants, which have been substantially addressed by mRNA vaccine platforms.

It has been shown that 2 doses of mRNA vaccine induced robust and durable antibody response lasting for 6∼9 months (37, 38). Different from the mRNA vaccine, nAb titers-elicited by the 2^nd^ CoronaVac vaccine waned rapidly from the peak levels. And most of the individuals (70/83) displayed no detectable nAb titers 6∼8 months after the 2^nd^ dose. Nevertheless, a 3^rd^ vaccine dose of CoronaVac vaccine significantly reinvigorated nAb responses. In particular, a 3^rd^ vaccine dose substantially improved the neutralization activities against Omicron B.1.1.529 and BA.2 variants. Of note, we found that the levels of SARS-CoV-2 IgG and nAb were negatively correlated with the age of participants at 2 weeks post the 3^rd^ dose, indicating the elder may poorly responded to the CoronaVac vaccine and a 4^th^ booster may be required.

In our longitudinal study from vaccinated individuals, we found that robust SARS-CoV-2 spike-specific memory CD4^+^ T cell responses following a 2^nd^ and 3^rd^ dose of CoronaVac vaccine in most of the participants. Moreover, we found that CoronaVac vaccination significantly altered the frequencies of polyclonal peripheral CD4^+^ T cell subsets, including a marked increase in the frequency of T_H_1 cells and the changes among cT_FH_ subsets. SARS-CoV-2 mRNA vaccines can induce robust antigen-specific T_FH_ responses in both peripheral blood and lymph nodes and maintained for 6 months (14, 15, 39). Similarly, our study found that CoronaVac vaccine efficiently elicited spike-specific, IFN-γ/IL-2 producing CD4^+^ T cells and cT_FH_ cells necessary for the anti-viral and antibody response (26, 40–42). And the expanded spike-specific cT_FH_ cells were biased to the proinflammatory cT_FH_17 subsets post the 2^nd^ and 3^rd^ dose of CoronaVac, which was also found after SARS-CoV-2 infection (43, 44). Interestingly, we found that circulating T_FH_1 cells were positively associated with effector CCR7^lo^PD1^hi^ cT_FH_ cells. These effector-memory like T_FH_ cells are known to indicate the T_FH_ cell activity in the secondary lymphoid organs and effectively respond to the vaccination (25). What’s new here is that we further found EM like CCR7^lo^PD1^hi^ proportion of T_FH_1 cells were particularly sensitive to the antigen and were rapidly boosted following the 2^nd^ and 3^rd^ CoronaVac vaccination. In contrast, cT_FH_17 cells were highly enriched with central-memory like CCR7^hi^PD1^lo^ cT_FH-CM_ cells. The frequency of cT_FH_17-CM cells remained stable over the time course of the administration of 3 vaccinations. Similar to the bulk cT_FH_ cells, the frequency of spike-specific cT_FH-EM_ cells were markedly invigorated by the the 2^nd^ and 3^rd^ dose of vaccine. These results further support the notion that cT_FH-EM_ cells may serve as a reliable biomarker when evaluating the effectiveness of vaccine induced humoral immune response. Targeting the cT_FH-EM_ cells may also improve the vaccine response, which is worthwhile investigating in future studies.

The positive correlation between cT_FH_1 cells and SARS-CoV-2 IgM, IgG titers was reported in COVID-19 convalescent individuals (24, 45) and in other infections (46–48). These evidence, coupled with our data herein, inspired us to interrogate whether the cT_FH_ cell subsets associated with spike-specific antibody response. Indeed, clear correlations between the polyclonal cT_FH_ cells and SARS-CoV-2 IgG and IgM antibody titers induced by a 3^rd^ dose of CoronaVac vaccine. And a significant positive correlation between polyclonal cT_FH_1 cells and IgM was also found. Interestingly, we also observed a positive correlation between the cell numbers of spike-specific cT_FH_ cells, cT_FH_1 cells and SARS-CoV-2 IgG and nAb titers two weeks after the 2^nd^ and 3^rd^ vaccination. In brief, our data manifested that CoronaVac induced T_FH_ responses were highly related to high-affinity antibody responses.

It should be noted that more striking relationship between the T_FH_ cell and humoral immune response were observed at T3 compared to T5. A possible explanation for this is the difference in sample size. We were able to collect more longitudinal samples at T3 than at T5, which may have resulted in a clearer statistical relationship at T3. Biologically speaking, at T3, vaccine recipients received a more classical “prime-boost” immunization where the germinal center response reaches its peak. In response to a foreign antigen, a robust T_FH_-GC B cell coordination is formed in a relatively clean system. While at T5, a more complex germinal center response was induced by a third vaccine boost, where a mixed GC response with a few long-lasting GCs and new GCs could be present in the same vaccinee. The phenomenon of “original antigenic sin” could also impact the recruitment of new B cell clones into the GC response after repeated exposure to the same antigen, and how this affects the T_FH_ cell response is unknown. It is highly likely that memory T_FH_ cells would compete with the new T_FH_ clones in providing help to the GC B cells, which may contribute to a less clear relationship between bulk T_FH_ cells and humoral immune response as the “help-kinetics” from memory T_FH_ cells and newly activated T_FH_ cells may be different after a third exposure to the same T-dependent antigen.

Sequential COVID-19 vaccinations with diverse vaccine platforms can effectively induce robust adaptive immune responses that provide protection against severe complications caused by SARS-CoV-2 and its subvariants (26, 40, 49). While the magnitudes of spike-specific and variant-specific antibody and T cell responses are mostly comparable between BNT162b2 and mRNA-1273, and higher than those induced by Ad26.COV2.S and NVX-CoV2373, direct comparison studies on comprehensive clinical and immunological parameters between inactivated COVID-19 vaccines such as CoronaVac and other vaccine platforms are limited (50–52). In general, inactivated vaccines like CoronaVac elicit relatively lower seropositivity and anti-spike RBD IgG antibody responses compared to mRNA vaccines like BNT162b2, and such antibody titers tend to wane faster than those induced by mRNA vaccines (53). Nevertheless, our study and others suggest that a third dose of CoronaVac can increase the overall and neutralizing antibody titers (Figure 1), potentially narrowing the quantitative gap of antibody titers between CoronaVac and mRNA vaccines (54). Interestingly, in line with some recent studies (55, 56), our data suggested that sequential administrations of CoronaVac induces robust effector and antigen-specific T cell response, similar to that of elicited by BNT162b2, mRNA-1273, Ad26.COV2.S, and protein-adjuvanted vaccine such as NVX-CoV2373 (50–52). Moreover, our data indicates that a 2^nd^ and 3^rd^ CoronaVac vaccination effectively boosts antigen-specific T_FH_ cells, the key helper T cells regulating antibody maturation, similar to mRNA vaccines (8, 51). Notably, we further identified that a CXCR3 expressing subset of T_FH_ cells, T_FH_1 cells represent the effector T_FH_ cells (T_FH-EM_), while T_FH_17 cells assemble the central memory like T_FH_ cells (T_FH-CM_) in response to sequential vaccinations. Further studies are needed to investigate whether mRNA and other vaccine types elicit a similar T_FH_ cell response.

There are several limitations in this study. First, the number of individuals enrolled after the 3^rd^ dose of CoronaVac is relatively small. This is in part due to some of the participates were infected by other virus such as influenza virus or vaccination by hepatitis B virus and no longer eligible based on our recruitment requirements. Our study, nevertheless, revealed the dynamics of polyclonal and SARS-CoV-2 specific CD4^+^ T cell response following each dose of CoronaVac vaccine. Despite the adequate knowledge acquired on this matter from mRNA-vaccinated individuals, there is a big knowledge gap in our understanding of T cell response elicited by inactivated SARS-CoV-2 vaccine, and how it may evolve longitudinally. Our study would provide necessary insights on this matter and guide the design of novel vaccine regimen of inactivated vaccine against SARS-CoV-2 virus and beyond.

## Acknowledgments

We are thankful to the donors for their blood samples used in this study.

## Author contributions

P.Zhou conceived and designed the study. C.C, T.J, Y.D and T.Z performed the experiments. F.G, Y.D, M.L and X.L participated in scientific discussion and recruited the patients. C.C, T.J, J.J, D.S and Z.B analyzed the data. C.C, T.J & F.G helped with the original illustration and draft. P.Z, C.C and T.J are co-first author and P.Z, X.L and F.G are co-corresponding author, order in which they are listed was determined by workload. P.Zhou wrote and revised the manuscript and led the submission. P.Zhou is the lead contact.

## Materials and methods

### Study design and Human subjects

A total of 88 participants (health care workers-HCW) who received 2 or 3 doses of SARS-CoV-2 vaccination (CoronaVac) at Affiliated Hospital of Jiangnan University and The Fifth People’s Hospital of Wuxi were recruited in our study from February 2021 and the study was initially done before December 2021. The medical ethical committee of the Affiliated Hospital of Jiangnan University (LS2021004) and The Fifth People’s Hospital of Wuxi (2020-034-1) reviewed and approved the study. Participants included were healthy adults aged 18 to 70 years without evidence of preceding SARS-CoV-2 infection. All individuals were nonatopic and with no infectious diseases or autoimmune diseases. Blood samples were collected at the following time points: pre-vaccination baseline (T1), 1 week after the 1^st^ dose (T2), 2 weeks after the 2^nd^ dose (T3), 6-8 months after the 2^nd^ dose (T4), as well as 2 weeks after the 3^rd^ dose (T5). The informed consent was obtained from all participants before sample collection.

### Peripheral blood mononuclear cells (PBMCs) and plasma isolation

Blood collection and processing were performed as previously described.(24) Briefly, whole blood was collected in EDTA-2K tubes (BD Biosciences) and processed for PBMCs and plasma isolation. EDTA-2K tubes were first centrifuged (450g, 5 min, 4°C), and the plasma was harvested for storage at −80 °C until required. Samples were further diluted with PBS (1:1) and separated using the Ficoll-Hypaque (GE Healthcare Life Sciences) density gradient (centrifugation 450g, 25 min, 20°C, without brake). PBMC layers were carefully collected and washed twice. After centrifugation, cells were resuspended in recovery media containing 10% dimethyl sulfoxide (Gibco), supplemented with 10% heat-inactivated fetal bovine serum (Gibco). Aliquots of cells were quickly transferred to freezing container (Corning Incorporated) at −80°C overnight. Samples were stored in liquid nitrogen until further use.

### Immunophenotyping by flow cytometry

Frozen aliquots of PBMCs were immediately thawed into prewarmed complete RPMI-1640 supplemented with 10% heat-inactivated FBS. Carefully washed once and resuspend in FACS buffer (PBS with 2% heat inactivated FBS). Add diluted 7-AAD (1:100 in the FACS buffer) to exclude dead cells and followed by Fc-receptor block (1:5 dilution, Miltenyi Biotec) to block non-specific staining. Cells were then stained with a cocktail of monoclonal antibodies including CD45RA-Alexa Fluor 488 (1:100, HI100), CD3-AF532 (1:200, UCHT1), CD4-Brilliant Violet 750 (1:200, SK3), CD8a-BV570 (1:200, RPA-T8), TCRγδ-BV480(1:100, B1), CD19-Super Bright 436 (1:100, HIB19), and CXCR5-APC (1:50, J252D4), CD25-PE (1:50, Clone: M-A251), CCR7-PE-Cy7 (CD197) (1:50, Clone: 3D12), HLA-DR-APC/Fire 750 (1:100, Clone: L243), CD183 (CXCR3)-BV421 (1:100, Clone: G025H7), CD196 (CCR6)-BV605 (1:100, Clone: G034E3), CD279 (PD-1)-BV650 (1:50, Clone: EH12.2H7), CD127-APC-R700 (1:100, Clone: HIL-7R-M21) **(Supplementary Table 1)**. After incubation for 30 min at 4°C in the dark, cells were washed twice in FACS buffer and then resuspended in 200 µL FACS buffer. Cells were kept on ice until acquisition.

Detailed immune phenotyping of CD4^+^ T cell using 24-color flow cytometry was performed on Cytek^TM^ Northern Lights with standardized configuration. Dead cells were routinely excluded from the analysis by staining with 7-AAD. For CD4^+^ T cells, EM (CD45RA^−^CCR7^−^), CM (CD45RA^−^CCR7^+^) and naïve cells (CD45RA^+^CCR7^+^) can be defined. Within the CD25 positive compartment, CD4^+^ T cells can be identified as T_REG_ (CD25^+^CD127^low^) and T_FR_ (CD25^+^CD127^low^CXCR5^+^PD-1^+^) cells. Within the CD25 negative compartment, CD4^+^ T cells can be divided into T_FH_ (CD45RA^−^CXCR5^+^), T_H_1 (CD45RA^−^CXCR5^−^CXCR3^+^CCR6^−^), T_H_2 (CD45RA^−^CXCR5^−^CXCR3^−^CCR6^−^), and T_H_17 (CD45RA^−^CXCR5^−^CXCR3^−^CCR6^+^). T_FH_ cells can be further divided into T_FH_1 (CXCR3^+^CCR6^−^), T_FH_2 (CXCR3^−^CCR6^−^) and T_FH_17 (CXCR3^−^CCR6^+^) cells. Data were analyzed with FlowJo software (Version 10).

### Activation-induced markers (AIM) T cell assay and intracellular staining (ICS) assay

Around 1×10^6^ cells per 200 μL were plated in 96-well U-bottom plate with complete RPMI-1640 medium containing 5% heat inactivated FBS. After resting overnight in the incubator at 37°C with 5% CO_2_, cells (1×10^6^) were stimulated with SARS-CoV-2 spike protein (S1+S2, 2 ug/mL, SinoBiological) in 5% CO_2_ at 37℃ for 24h. Co-stimulatory anti-CD28(1 ug/mL, Biolegend) and anti-CD49d (1 ug/mL, Biolegend) were added. SEB (1 ug/mL, Toxin Technology) were used as positive control and an equimolar amount of DMSO as negative control. Antigen-specific CD4^+^ T (HLA-DR^+^CD25^+^), and antigen-specific T_FH_ cells (CXCR5^+^ HLA-DR^+^CD25^+^) were defined by the AIM assay.

For ICS assay, 1×10^6^ PBMCs were cultured in the presence of SARS-CoV-2 spike peptide pools (S1+S2, 2 ug/mL, SinoBiological) for 24 hours at 37°C. Co-stimulatory anti-CD28(1 ug/mL, Biolegend) and anti-CD49d (1 ug/mL, Biolegend) were added. In addition, SEB (1 ug/mL, Toxin Technology) was used as positive control and an equimolar amount of DMSO was used as negative control. After 24 hours, Brefeldin A (1:1000, eBioscience) was added to the culture for an additional 4 hours. After incubation, cells were washed and stained with Fixable Viability Dyes (eFluor 520, eBioscience) for 30 minutes at 4°C. Cells were then stained with a cocktail of monoclonal antibodies for cell surface staining with Fc block. After surface staining, cells were permeabilized and stained with intracellular antibodies against IFN-γ-PE-Cy7 (1:50, Clone: 4S.B3) and IL-2-APC-R700 (1:50, Clone: MQ1-17H12) for 30 minutes in dark at room temperature. Cells were kept on ice until acquisition by Cytek^TM^ Northern Lights flow cytometer.

### SARS-CoV-2 IgG/IgM/IgA antibodies measurement

The concentrations of plasma SARS-CoV-2 IgG (Autobio Diagnostics), IgM (Autobio Diagnostics), and IgA (Beijing Wantai Biolodical) were measured by chemiluminescent microparticle immunoassay kits, according to the manufacturer’s instructions. The assay is based upon the two-step indirect method. Briefly, SARS-CoV-2 IgG/IgM/IgA present in the sample binds to the SARS-CoV-2 antigen coated microparticles. Then, HRP-conjugated anti-human IgG/IgM/IgA and followed by a chemiluminescent substrate was added into the reaction system, resulting in a chemiluminescent reaction. The resulting chemiluminescent reaction is measured as relative light unit (RLU), which is proportional to the amount of SARS-CoV-2 IgG/IgM/IgA in the samples. Results were evaluated by sample RLU/cut-off value (S/CO). Samples with S/CO values <1.00 are considered nonreactive (NR). Samples with S/CO values ≥1.00 are considered reactive (R).

### SARS-CoV-2 neutralizing antibodies measurement

The concentrations of plasma SARS-CoV-2 neutralizing antibodies were tested by chemiluminescent microparticle immunoassay (Autobio Diagnostics), according to the manufacturer’s instructions. This assay is based upon the one-step competitive method. The amount of SARS-CoV-2 neutralization antibodies in the samples is measured from the RLU by means of the stored calibration data and determined automatically by the system software. Samples with values <30 AU/mL are NR, values ≥ 30 AU/mL are R.

### SARS-CoV-2 Pseudovirus (PSV) neutralization assay

Pseudotyped HIV virus incorporated in different variants of SARS-CoV-2 S proteins (Vazyme) were used to test the neutralizing activity of serum from vaccination recruits. The SARS-CoV-2 pseudoviruses bearing wild-type, B.1.1.529, BA.2 spike protein were provided by Vazyme. Serum samples were first heat-inactivated in a water bath for 30 min at 56 °C and then serially diluted in three-folds with complete DMEM medium from 1:20 to 1:4860 in 96-well flat-bottom culture plates in a total volume of 150 μL. The cell control (CC) with only cells and the virus control (VC) with virus and cells were set up in each plate. The SARS-CoV-2 pseudotyped virus were diluted to 2 × 10^4^ TCID_50_/mL in complete DMEM, and 50 μL diluted pseudotyped virus were added to each well and incubated for 1 h at 37 °C. The sample wells were finally diluted from 1:30 to 1:7290. Adjust the HEK293-ACE2 (Vazyme) cell concentration to 4 × 10^5^ cells/mL with DMEM complete medium and add 50 μL of cell suspension into all wells and incubated for 48h at 37 °C and 5% CO_2_. Finally, Bio-Lite Luciferase Assay System (Vazyme) was employed to measure the firefly luciferase activity, to obtain the neutralizing antibody content of the sample. Neutralizing antibody titers were calculated as ID_50_ expressed as the dilution of serum that resulted in a 50% reduction of luciferase luminescence compared with a virus control.

## QUANTIFICATION AND STATISTICAL ANALYSIS

Concentrations of SARS-CoV-2 anti-spike IgG, anti-spike IgM, anti-spike IgA, and neutralizing antibodies in vaccinated individuals between five different time points were compared using Wilcoxon matched-pairs signed-ranks test. The non-parametric Mann-Whitney U test was used to compare the effects of different time intervals between the 1^st^ and 2^nd^ dose on SARS-CoV-2 antibody levels. The frequency of polyclonal peripheral CD4^+^ T and spike-specific CD4^+^ T cell subsets were calculated using Wilcoxon matched-pairs signed-rank test for comparison between different time points. The two-tailed, non-parametric Spearman’s rank correlation test and Pearson test were used to evaluate the correlations between CD4^+^ T cells and antibody responses following CoronaVac vaccination, and the correlation between age and SARS-CoV-2 antibody titers. Statistical analysis was carried out using GraphPad Prism (V 9.2.0) software, and the correlation matrix mapping used R (V 3.6.3) software. *P* values are indicated with asterisks: \**P* < 0.05; \*\**P* < 0.01; *** *P* < 0.001; **** *P* < 0.0001.

---

**Supplementary table 1.**
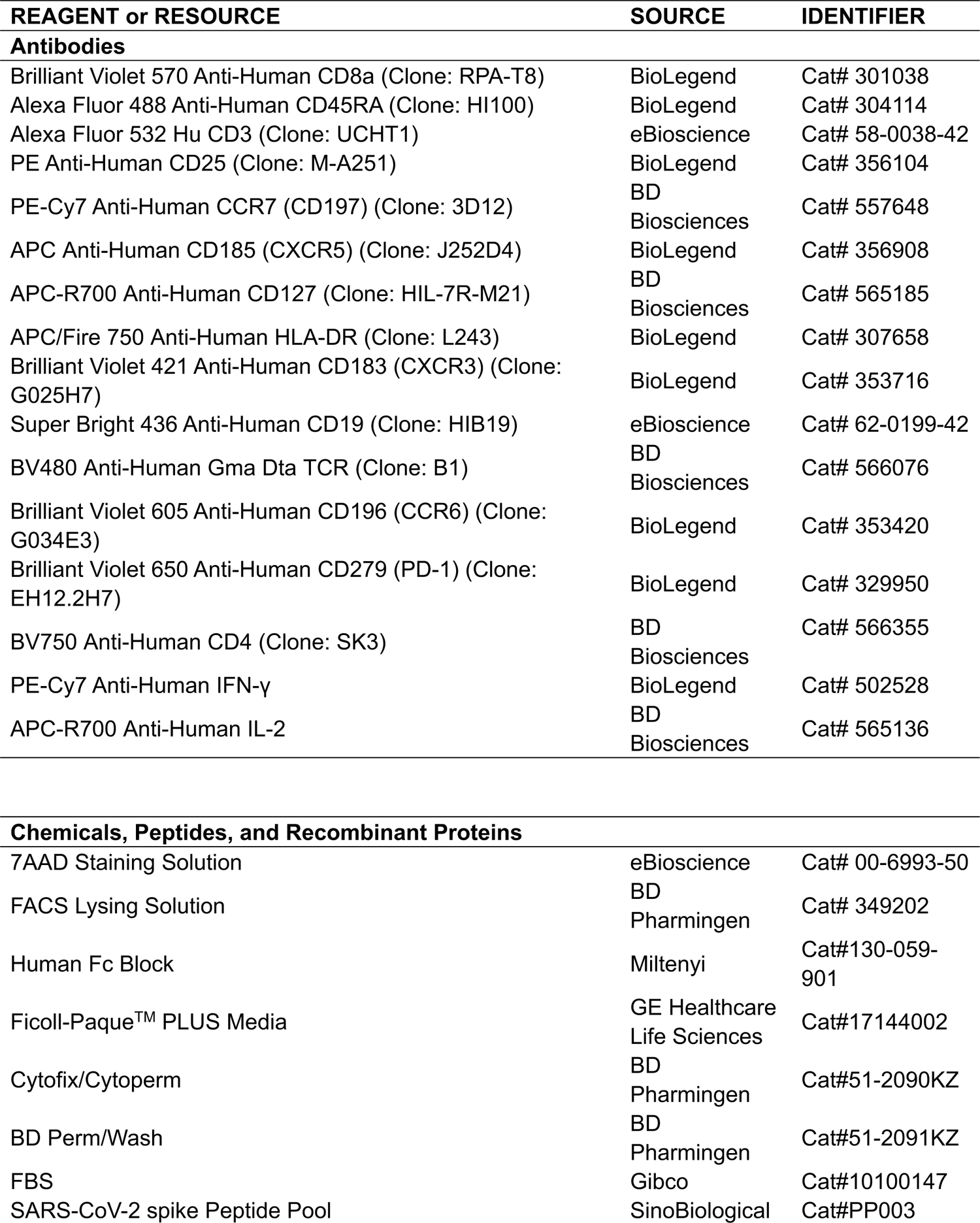
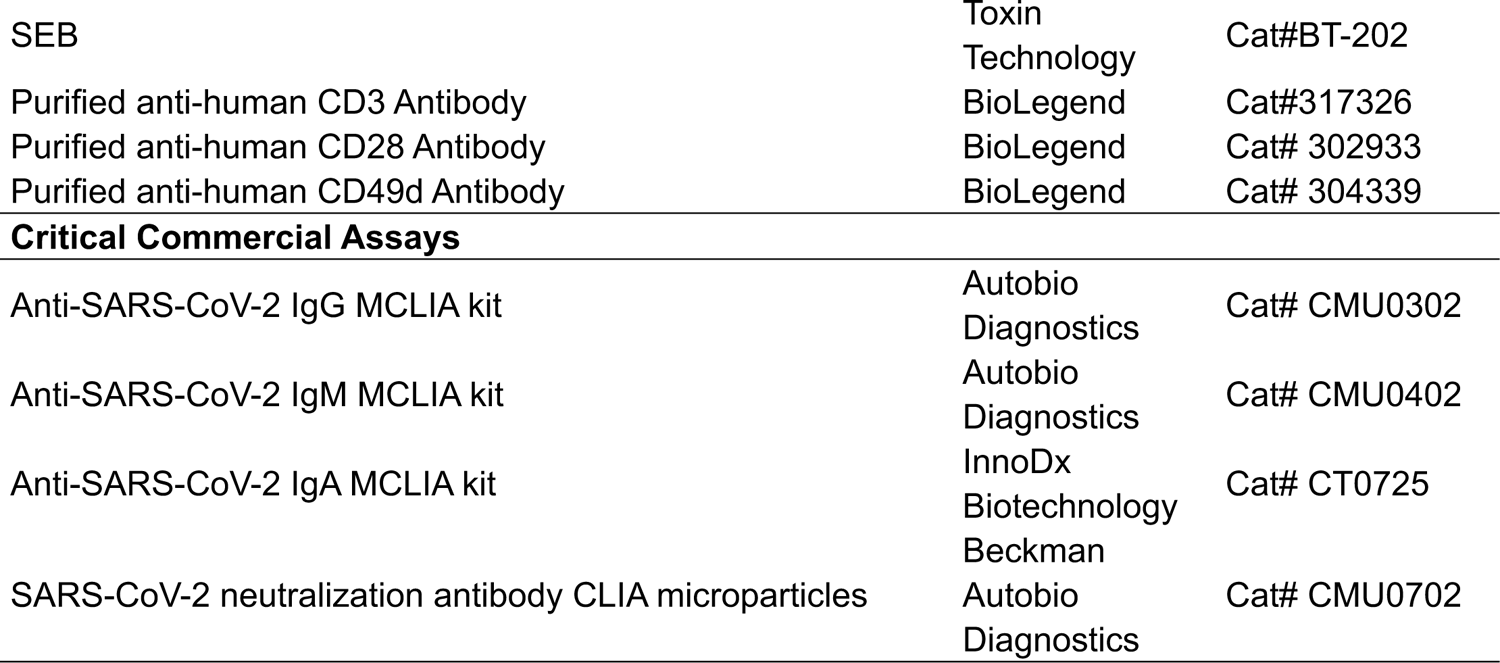
Antibodies and other key resources

