## Supplementary figure for "Longitudinal analysis of memory T follicular helper cells and antibody response following CoronaVac vaccination"

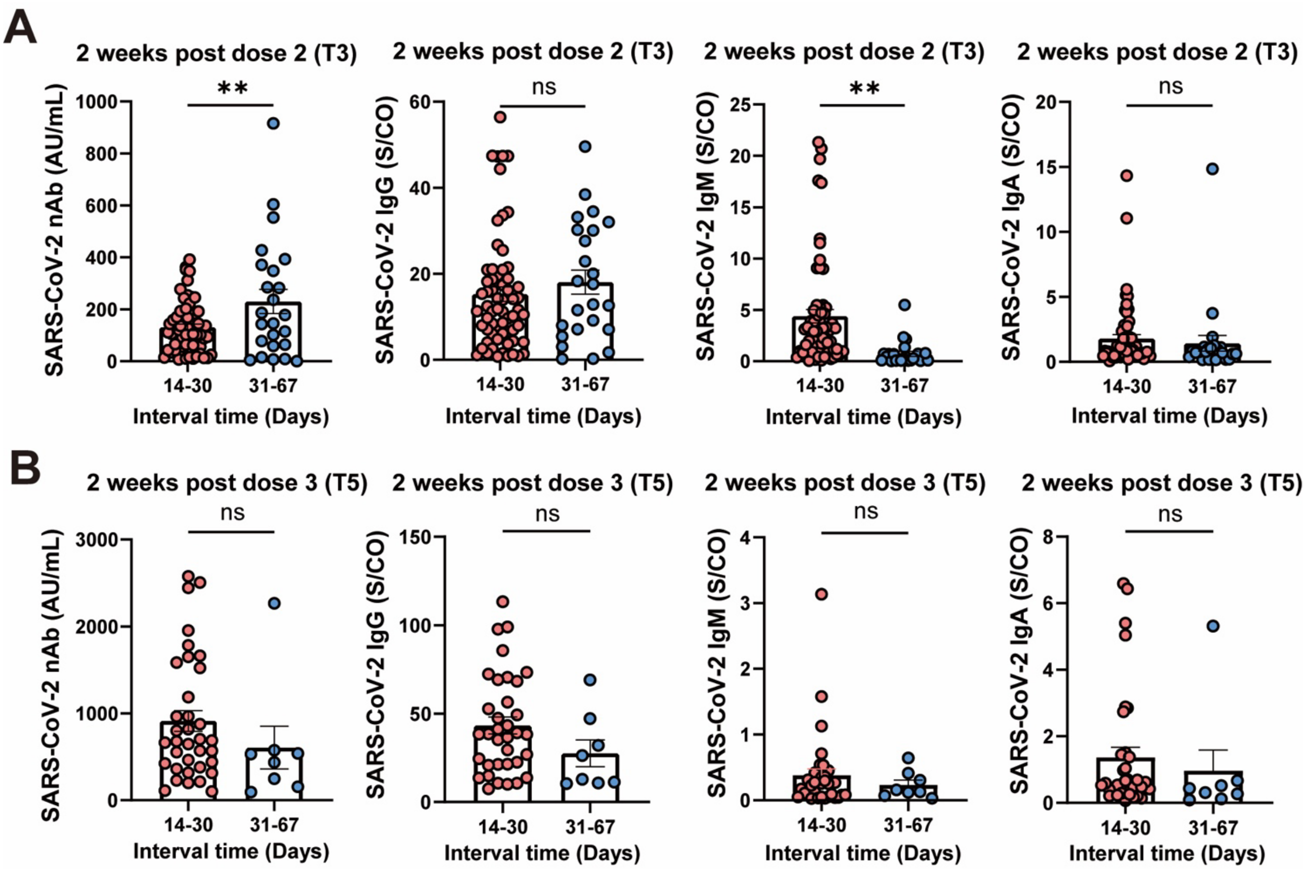


**Supplementary figure S1: Vaccination interval and the levels of SARS-CoV-2-specific antibodies induced by CoronaVac vaccine**

Effect of different time intervals between the primary and second dose on the levels of SARS-CoV-2 IgM, IgG, IgA and nAb at 2 weeks after the 2$^{nd}$ dose of vaccine (T3) (A) and 2 weeks after the 3$^{rd}$ dose of vaccine (T5) (B). Each dot represents an individual subject. Bars represent the mean values with SEM. Statistics were calculated using Mann-Whitney U test (A, B). *$P$ < 0.05; **$P$ < 0.01; ns, not significant.

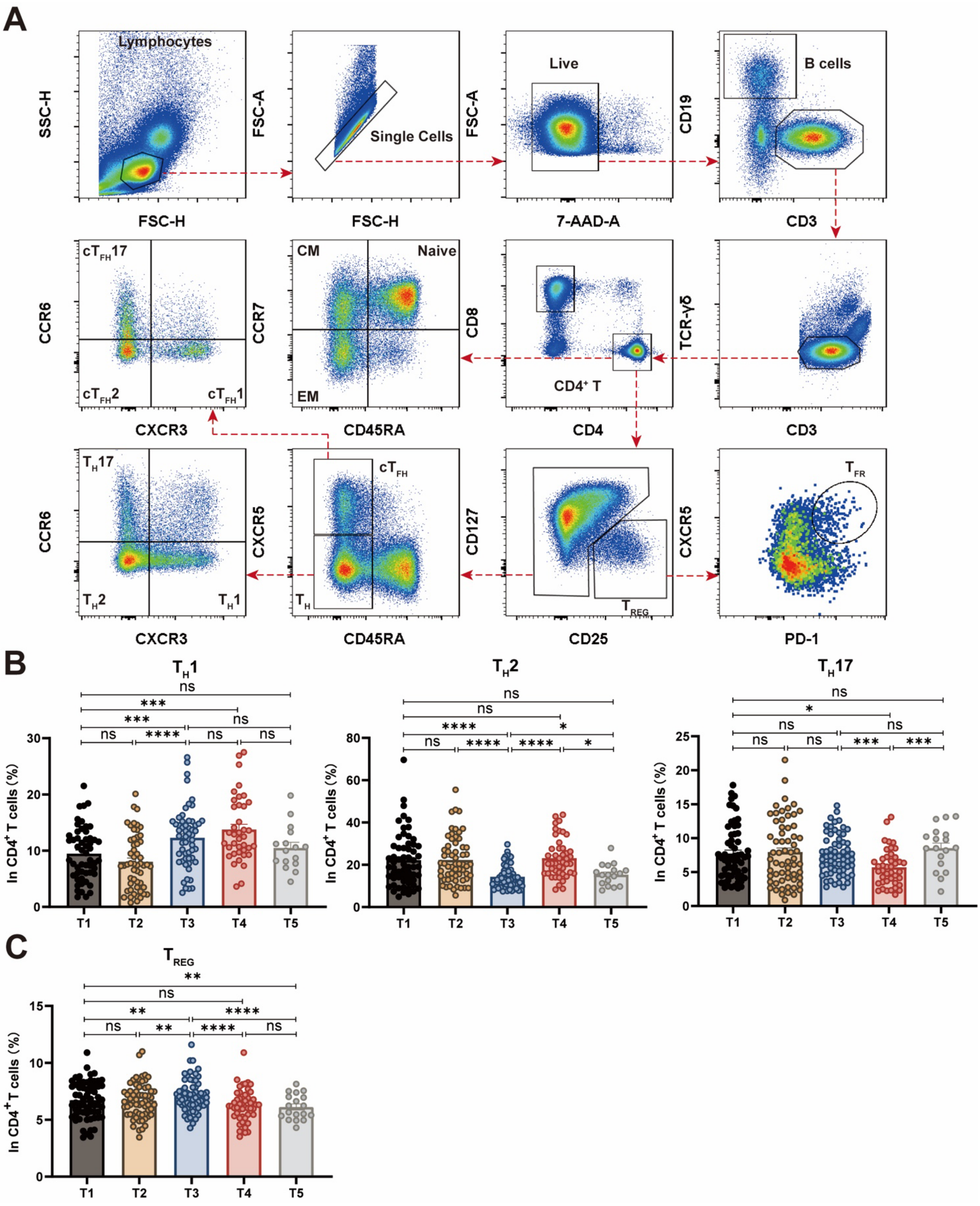


**Supplementary figure S2: Polyclonal circulating CD4⁺ T cells.**

(A) Representative plots illustrate the gating strategy of polyclonal CD4⁺ T cells. Lymphocytes were firstly gated, and doublets were excluded by FSC-H and FSC-A signals. Live cells were identified by 7-AAD. $CD4^+$ T cells were gated on $CD3^+CD19^-TCR\gamma\delta^-$ cells. In particular, EM ($CD45RA^-CCR7^-$), CM ($CD45RA^-CCR7^+$) and naïve cells ($CD45RA^+CCR7^+$) are defined accordingly. $T_{REG}$ ($CD25^+CD127^{low}$) and $T_{FR}$ ($CD25^+CD127^{low}CXCR5^+PD\text{-}1^+$) cells are gated accordingly. CD25 negative cells can be divided into $cT_{FH}$ ($CD45RA^-CXCR5^+$) cells and $T_H$ ($CD45RA^-CXCR5^-$) cells. Based on the expression of CXCR3 and CCR6, $cT_{FH}$ cells were further divided into $cT_{FH}1$ ($CXCR3^+CCR6^-$), $cT_{FH}2$ ($CXCR3^-CCR6^-$) and $cT_{FH}17$ ($CXCR3^-CCR6^+$) cells, while $T_H$ cells were divided into $T_H1$ ($CXCR3^+CCR6^-$), $T_H2$ ($CXCR3^-CCR6^-$), and $T_H17$ ($CXCR3^-CCR6^+$) cells. Longitudinal dynamics of polyclonal $T_H1$, $T_H2$, $T_H17$ (B) and $T_{REG}$ (C) cells during three doses of vaccines at five timepoints. Each dot represents an individual subject. Bars represent the mean values with SEM. Statistics were calculated using Wilcoxon matched-pairs signed rank for comparison between timepoints (B, C). *, $P < 0.05$; **, $P < 0.01$; ***, $P < 0.001$; ****, $P < 0.0001$; ns, not significant.

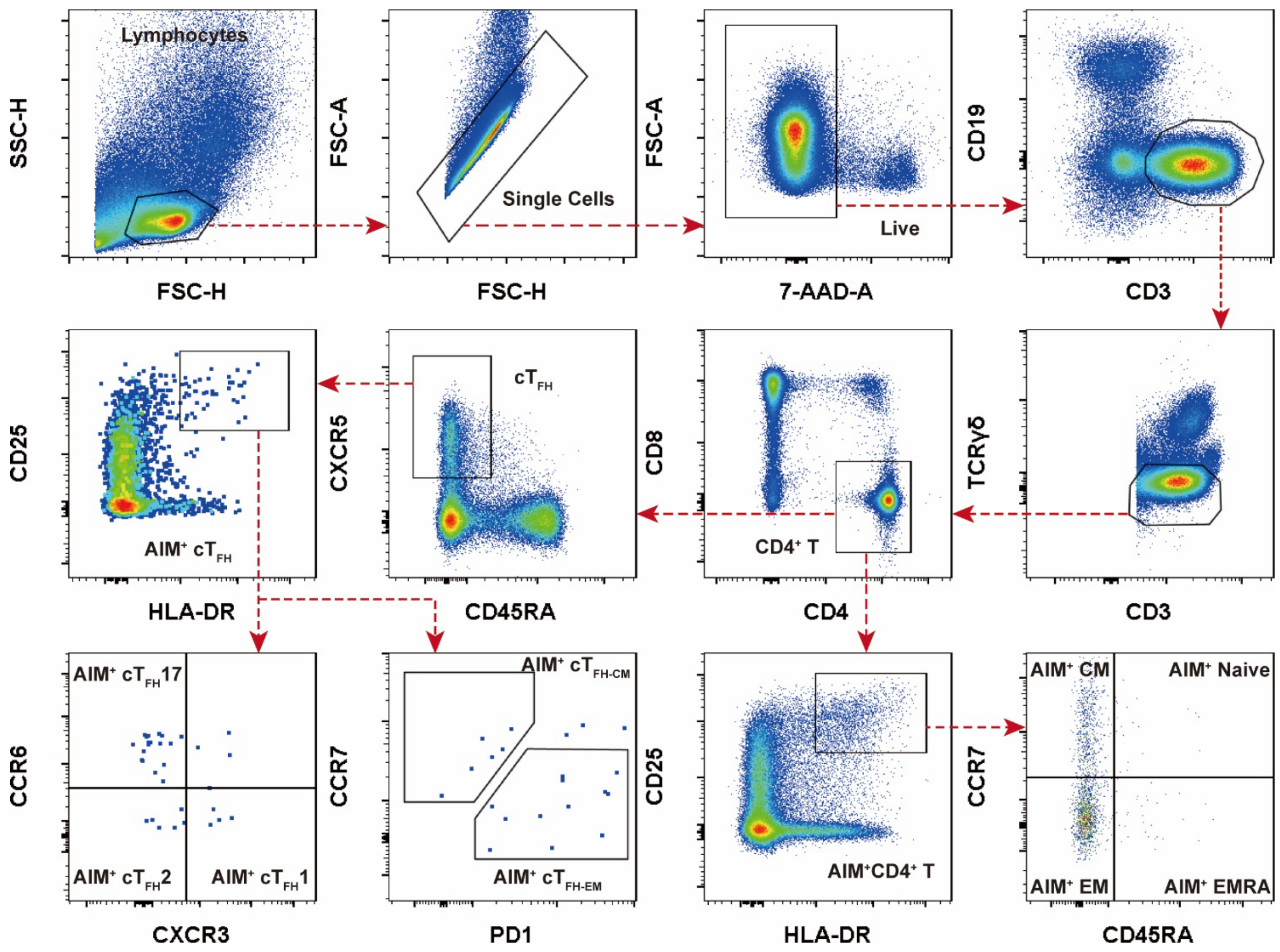


**Supplementary figure S3: Gating-strategy for spike-specific CD4⁺ T and cT$_{FH}$ cells.** PBMCs were stimulated with SARS-CoV-2 spike protein (S1+S2, 2 ug/mL, SinoBiological) for 24h. Representative plots illustrate the gating strategy. CD4$^+$ T cells were driven from CD3$^+$CD19$^-$TCRγδ$^-$ gating. Spike-specific CD4$^+$ T cells and spike-specific cT$_{FH}$ cells were defined as HLA-DR$^+$CD25$^+$CD4$^+$ and CXCR5$^+$ HLA-DR$^+$CD25$^+$CD4$^+$, respectively. Spike-specific CD4$^+$ T cells were further divided into memory and naïve cells. Spike-specific cT$_{FH}$ cells were further divided into cT$_{FH}$1, cT$_{FH}$2 and cT$_{FH}$17 cells according to the expression of surface molecules CXCR3 and CCR6, and divided into AIM$^+$ cT$_{FH-CM}$ and cT$_{FH-EM}$ cells by CCR7 and PD1.

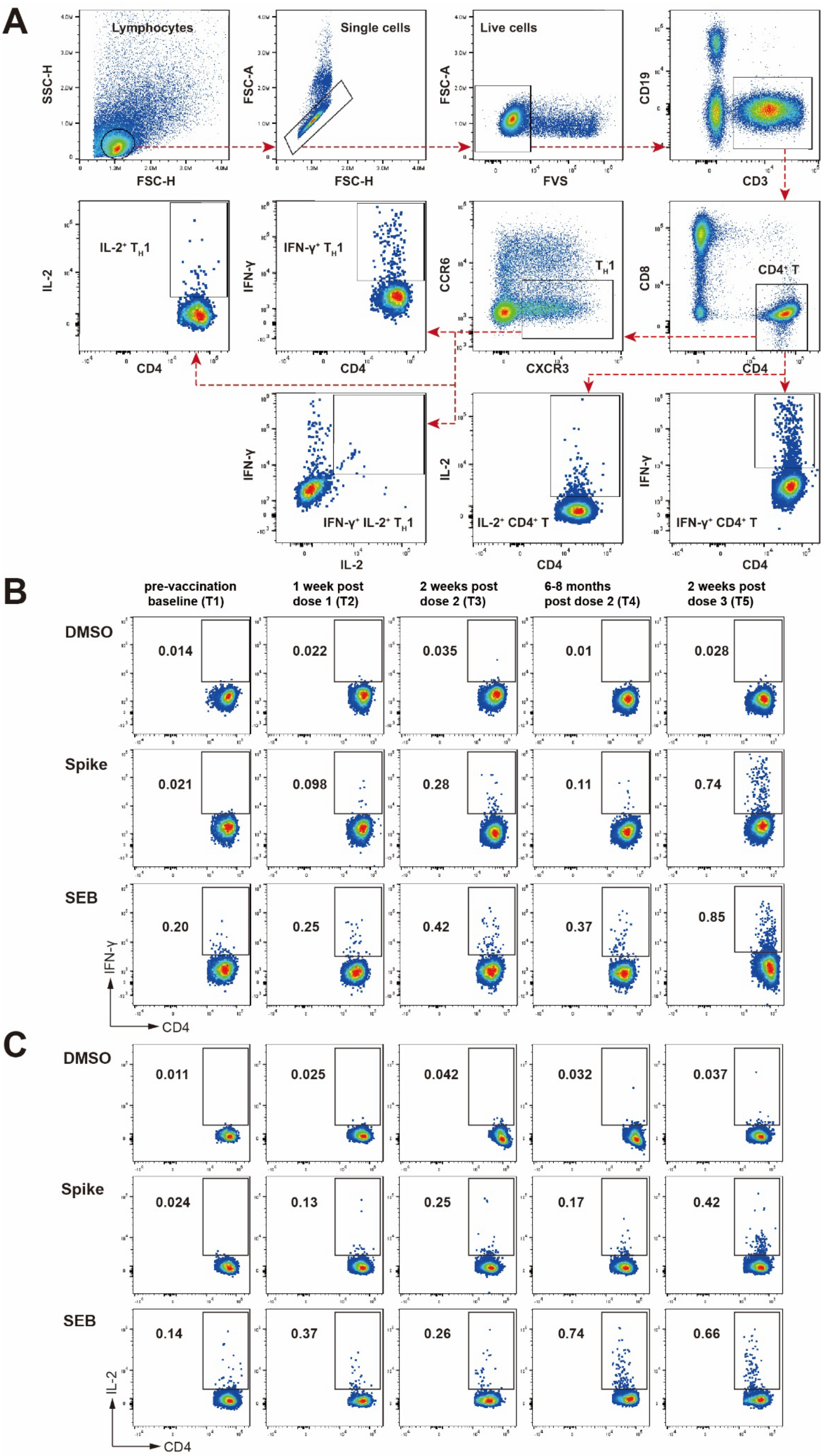

A
Lymphocytes
SSC-H
FSC-H
Single cells
FSC-A
Live cells
FVS
CD19
CD3
IL-2⁺ T_H1
IL-2
CD4
IFN-γ⁺ T_H1
IFN-γ
CCR6
CXCR3
T_H1
CD8
CD4⁺ T
IFN-γ⁺ IL-2⁺ T_H1
IL-2⁺ CD4⁺ T
IFN-γ⁺ CD4⁺ T
B
pre-vaccination baseline (T1)
1 week post dose 1 (T2)
2 weeks post dose 2 (T3)
6-8 months post dose 2 (T4)
2 weeks post dose 3 (T5)
DMSO
0.014
0.022
0.035
0.01
0.028
Spike
0.021
0.098
0.28
0.11
0.74
SEB
0.20
0.25
0.42
0.37
0.85
IFN-γ
CD4
C
DMSO
0.011
0.025
0.042
0.032
0.037
Spike
0.024
0.13
0.25
0.17
0.42
SEB
0.14
0.37
0.26
0.74
0.66
IL-2
CD4

**Supplementary figure S4: Gating-strategy and representative plots for spike-specific $CD4^+$ T cells using ICS assay.**

(A) Representative plots illustrate the gating strategy of spike-specific $CD4^+$ T cells. $CD4^+$ T cells were gating from live $CD3^+CD19^-$ lymphocytes. $T_H1$ cells were defined as $CXCR3^+$ $CCR6^-$ $CD4^+$ T cells. IFN-$\gamma^+$ $CD4^+$ T cells, IL-$2^+$ $CD4^+$ T cells, IFN-$\gamma^+$ $T_H1$, IL-$2^+$ $T_H1$ and IFN-$\gamma^+$ IL-$2^+$ $T_H1$ were further gated based on the cytokine production. Representative FACS plots of IFN-$\gamma^+$ $CD4^+$ T cells (B) and IL-$2^+$ $CD4^+$ T cells (C) were shown at five time points (T1-T5). DMSO and SEB were used as negative and positive controls, respectively.

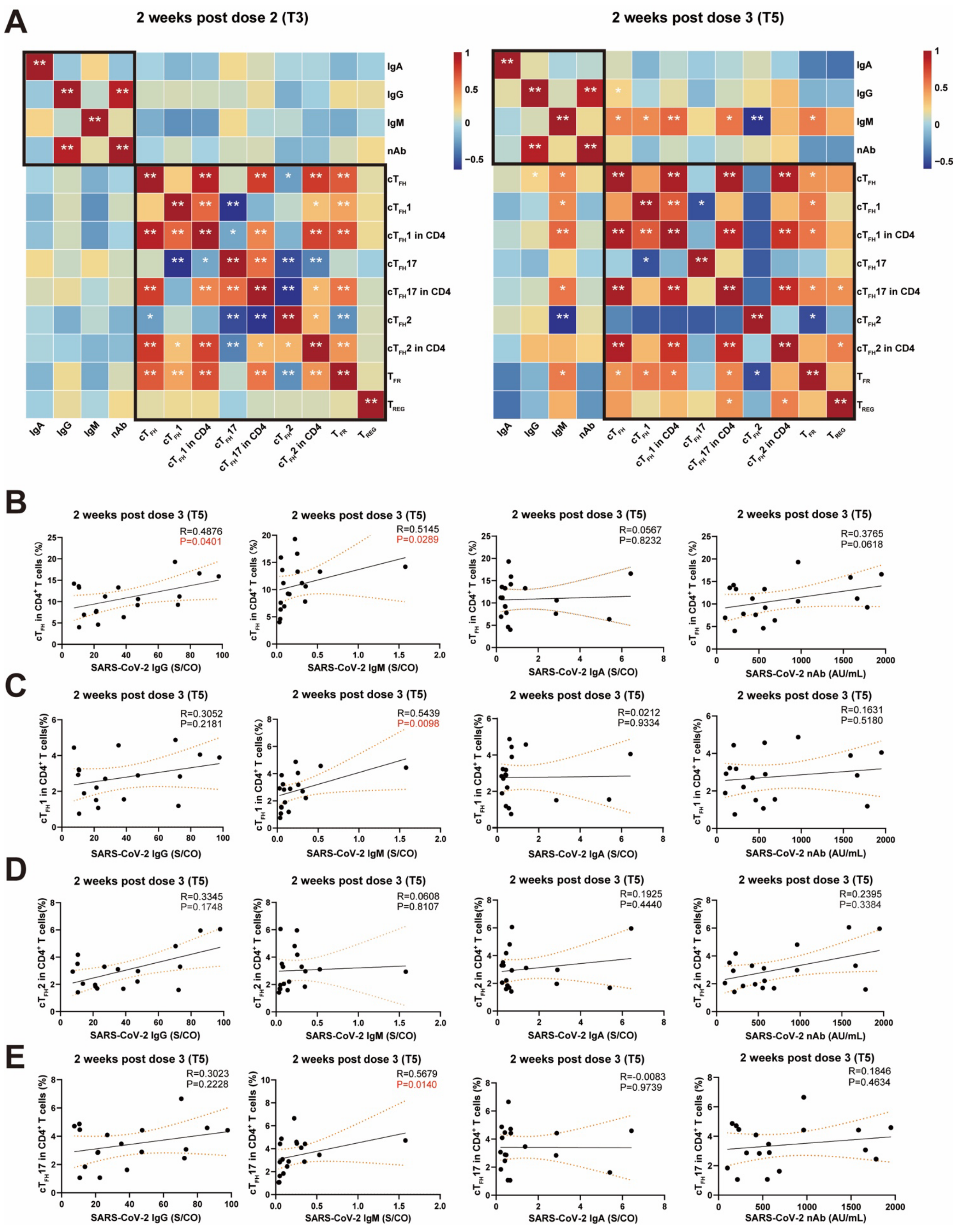

**Supplementary figure S5: Correlations between polyclonal CD4$^+$ T cells and antibody responses following CoronaVac vaccination.**

(A) Correlation heatmaps of the polyclonal $CD4^+$ T cell subsets and SARS-CoV-2 specific antibodies at two weeks post dose 2 (T3) and dose 3 (T5). Correlation analysis between the frequency of polyclonal $cT_{FH}$ cells (B), $cT_{FH}1$ cells (C), $cT_{FH}2$ cells (D), $cT_{FH}17$ cells (E) and SARS-CoV-2-specific IgG, IgM, IgA and nAb titers at two weeks post dose 3 (T5). Each dot represents an individual subject. The non-parametric Spearman's rank correlation was used (A, B, C, D, E). $^*P < 0.05$; $^{**}P < 0.01$ (A). *P* and R values were indicated (B, C, D, E).
